# Id proteins promote a cancer stem cell phenotype in triple negative breast cancer via negative regulation of Robo1

**DOI:** 10.1101/497313

**Authors:** Wee S. Teo, Holly Holliday, Nitheesh Karthikeyan, Aurélie S. Cazet, Daniel L. Roden, Kate Harvey, Christina Valbirk Konrad, Reshma Murali, Binitha Anu Varghese, Archana P. T., Chia-Ling Chan, Andrea McFarland, Simon Junankar, Sunny Ye, Jessica Yang, Iva Nikolic, Jaynish S. Shah, Laura A. Baker, Ewan K.A. Millar, Mathew J. Naylor, Christopher J. Ormandy, Sunil R. Lakhani, Warren Kaplan, Albert S. Mellick, Sandra A. O’Toole, Alexander Swarbrick, Radhika Nair

## Abstract

Breast cancers display phenotypic and functional heterogeneity and several lines of evidence support the existence of cancer stem cells (CSCs) in certain breast cancers, a minor population of cells capable of tumor initiation and metastatic dissemination. Identifying factors that regulate the CSC phenotype is therefore important for developing strategies to treat metastatic disease. The Inhibitor of Differentiation Protein 1 (Id1) and its closely related family member Inhibitor of Differentiation 3 (Id3) (collectively termed Id) are expressed by a diversity of stem cells and are required for metastatic dissemination in experimental models of breast cancer. In this study, we show that ID1 is expressed in rare neoplastic cells within ER-negative breast cancers. To address the function of Id1 expressing cells within tumors, we developed two independent murine models of Triple Negative Breast Cancer (TNBC) in which a genetic reporter permitted the prospective isolation of Id1^+^ cells. Id1^+^ cells are enriched for self-renewal in tumorsphere assays *in vitro* and for tumor initiation *in vivo*. Conversely, depletion of Id1 and Id3 in the 4T1 murine model of TNBC demonstrates that Id1/3 are required for cell proliferation and self-renewal *in vitro*, as well as primary tumor growth and metastatic colonization of the lung *in vivo*. Using combined bioinformatic analysis, we have defined a novel mechanism of Id protein function via negative regulation of the Roundabout Axon Guidance Receptor Homolog 1 (*Robo1*) leading to activation of a Myc transcriptional programme.

## Introduction

Several lines of evidence suggest that rare sub-populations of tumor cells, commonly termed cancer stem cells (CSCs), drive key tumor phenotypes such as self-renewal, drug resistance and metastasis and contribute to disease relapse and associated patient mortality (Chen et al., 2012; Lawson et al., 2015; Li et al., 2008; Malanchi et al., 2011). Recent evidence points to the hypothesis that CSCs are not static, but they exist in dynamic states, driven by critical transcription factors and are highly dependent on the microenvironmental cues (da Silva-Diz et al., 2018; Lee et al., 2016b; Wahl and Spike, 2017). Understanding the molecular networks that are critical to the survival and plasticity of CSCs is fundamental to resolving clinical problems associated with chemo-resistance and metastatic residual disease.

The Inhibitor of DNA binding (ID) proteins have previously been recognized as regulators of CSCs and tumor progression (Lasorella et al., 2014). These proteins constitute a family of four highly conserved transcriptional regulators (ID1-4) that act as dominant-negative inhibitors of basic helix–loop–helix (bHLH) transcription factors. ID proteins are expressed in a tissue-specific and stage-dependent manner and are required for the maintenance of self-renewal and multipotency of embryonic and many tissue stem cells (Aloia et al., 2015; Hong et al., 2011; Liang et al., 2009; Stankic et al., 2013). Previous studies have reported a functional redundancy among the four members of the mammalian Id family, in particular Id1 and Id3 (referred to collectively here as Id), and their overlapping expression patterns during normal development and cancer (Anido et al., 2010; Gupta et al., 2007; Lyden et al., 1999; Niola et al., 2013; O’Brien et al., 2012). Id2 and Id4 were not investigated in this work as they are found to have independent functions from Id1 and Id3.

A number of studies have implied a significant role for ID1 and ID3 in breast cancer progression and metastasis (Gupta et al., 2007). We have previously demonstrated that Id1 cooperates with activated Ras signalling and promotes mammary tumor initiation and metastasis *in vivo* by supporting long-term self-renewal and proliferative capacity (Swarbrick et al., 2008). Additional work has clearly implicated ID1 in regulating D- and E-type cyclins and their associated cyclin-dependant kinases, CDK4 and CDK2 in human breast epithelial cells, p21 (Swarbrick et al., 2005), the matrix metalloproteinase MT1-MMP (Fong et al., 2003), KLF17 (Gumireddy et al., 2009), Cyclin D1 (Tobin et al., 2011), Bcl-2 (Kim et al., 2008), and BMI1 (Qian et al., 2010) among others.

Even though several Id-dependent targets have been identified, we still lack a comprehensive picture of the downstream molecular mechanisms controlled by Id and their associated pathways mediating breast cancer progression and metastasis particularly in the poor prognostic TNBC subtype. In this study, we demonstrate using four independent mouse models of TNBC that Id is important for the maintenance of a CSC phenotype. We also describe a novel mechanism by which Id controls the CSC state by negatively regulating Robo1 to control proliferation and self-renewal via indirect activation of a Myc transcriptional programme.

## Materials and Methods

### Plasmids

pEN_TmiRc3 parental entry plasmid, pSLIK-Venus and pSLIK-Neo destination vectors were obtained from the ATCC (Manassas, VA,USA).

### Cell culture

4T1 and HEK293T cells were obtained from the American Type Culture Collection (ATCC). 4T1 cells were maintained in RPMI 1640 (Gibco, Grand Island, NY, USA) supplemented with 10% (v/v) FBS (Thermo Fisher Scientific, Scoresby, Vic, Australia), 20mM HEPES (Gibco, Grand Island, NY, USA), 1mM sodium pyruvate (Gibco, Grand Island, NY, USA), and 0.25% (v/v) glucose. HEK293T cells were grown in DMEM (Gibco, Grand Island, NY, USA) supplemented with 10% (v/v) FBS (Thermo Fisher Scientific, Scoresby, Vic, Australia), 6mM L-glutamine (Gibco, Grand Island, NY, USA), 1mM sodium pyruvate (Gibco, Grand Island, NY, USA) and 1% (v/v) MEM Non-essential Amino Acids (Gibco, Grand Island, NY, USA). All cell lines were cultured at 37°C in a humidified incubator with 5% CO2.

### Animals

All experiments involving animal work were performed in accordance with the rules and regulations stated by the Garvan Institute Animal Ethics Committee. The BALB/c mice were sourced from the Australian BioResources Ltd. (Moss Vale, NSW, Australia). FVBN mice, p53 null mice, C3-Tag mice were a generous gift from Tyler Jacks, Cambridge, MA. Doxycycline (Dox) food, which contains 700mg Dox/kg, was manufactured by Gordon’s Specialty Stock Feed (Yanderra, NSW, Australia) and fed to the mice during studies involving Dox-induced knockdown of Id1/3.

### mRNA and Protein expression analysis

Total RNA from the cells were isolated using Qiagen RNeasy minikit (Qiagen, Doncaster, VIC, Australia) and cDNA was generated from 500 ng of RNA using the Superscript III first strand synthesis system (Invitrogen, Mulgrave, VIC, Australia) according to the manufacturer’s protocol. Quantitative real-time PCR was carried out using the TaqMan probe-based system (Applied Biosystems/Life Technologies, Scoresby, Vic, Australia) on the ABI Prism 7900HT Sequence Detection System (Biosystems/Life Technologies, Scoresby, Vic, Australia) according to manufacturer’s instructions. The probes used for the gene expression analysis by TaqMan assay are; Mouse Id1-Mm00775963_g1, Mouse Id3-Mm01188138_g1, Mouse Robo1-Mm00803879_m1, Mouse Fermt1-Mm01270148_m1, Mouse Foxc2-Mm00546194_s1, Mouse Gapdh-Mm99999915_g1 and Mouse β-Actin-Mm00607939_s1. For protein expression analysis, lysates were prepared in RIPA lysis buffer supplemented with complete ULTRA protease inhibitor cocktail tablets (Roche, Basel, Switzerland) and western blotting was performed as demonstrated before (Nair et al., 2014a). The list of antibodies used for western blotting are given in Supplementary Table S6.

### Immunohistochemistry

Immunohistochemistry analysis was performed as described earlier (Nair et al., 2014a). Briefly, 4µm-thick sections of formalin-fixed, paraffin-embedded (FFPE) tissue blocks were antigen retrieved by heat-induced antigen retrieval and were incubated with respective primary and secondary antibodies (listed in Supplementary Table S7).

### Id1GFP reporter in the p53-/- model

p53-/- tumors arise spontaneously following transplantation of Tp53-null mammary epithelium into the mammary fat pads of naïve FVB/n mice. The tumors were then transplanted into naïve recipients; this method has been previously used to study murine TNBC CSCs (Herschkowitz et al., 2007; Hochgrafe et al., 2010). We developed and validated an Id1/GFP molecular reporter construct in which 1.2kb of the Id1 proximal promoter is placed upstream of the GFP cDNA. Cells with active Id1 promoter can be visualized and isolated based on GFP expression by FACS from primary mouse tumors and cell lines. A similar approach has been successfully used to isolate CSCs with active β-catenin signalling (Zhang et al., 2010). Using the reporter construct, we typically see between 2-15% of cancers cells are GFP^+^ by FACS, depending on the clone analysed. We experimentally validated the Id1/GFP system to ensure that GFP expression accurately marks the Id1 cells within the bulk tumor cell population. After transfection of the Id1/GFP reporter into cultured p53-/- tumor cells, both the sorted GFP^+^ and unsorted cells were able to generate new tumors when transplanted into wild-type recipient mice. Tumors were harvested, dissociated into single cells, expanded briefly *in vitro*, and then FACS sorted once more to collect GFP^+^ and GFP+ cell fractions.

### Generation of shRNA lentiviral vectors

Single stranded cDNA sequences of mouse Id1 and Id3 shRNAs were purchased from Sigma-Aldrich (Lismore, NSW, Australia). The Id1 shRNA sequence which targets 5’-GGGACCTGCAGCTGGAGCTGAA-3’ has been validated earlier (Gao et al., 2008). The Id3 sequence was adopted from Gupta et al 2007 (Gupta et al., 2007) and targets the sequence 5’-ATGGATGAGCTTCGATCTTAA-3’.shRNA directed against EGFP was used as the control. The shRNA linkers were designed as described earlier (Shin et al., 2006). The sense and antisense oligonucleotides with BfuAI restriction overhangs were annealed and cloned into the BfuAI restriction siteofpEN_TmiRc3 entry plasmid. pSLIK lentiviral vectors expressing shRNA against Id1 and Id3namely pSLIK-Venus-TmiR-shId1 and pSLIK-Neo-TmiR-shId3,were generated by Gateway recombination between the pEN_TmiR_Id1 or the pEN_TmiR_Id3 entry vector and the pSLIK-Venus or pSLIK-Neo destination vector respectively. Control pSLIK vector expressing shRNA against EGFP (pSLIK-Neo-TmiR-shEGFP) was generated by recombination between the pEN_TmiR_EGFP vector and the pSLIK-Neo vector. The Gateway recombination was performed using the LR reaction according to the manufacturer’s protocol (Invitrogen, Mulgrave, Vic, Australia).

### Lentivirus production

Lentiviral supernatant was produced by transfecting each lentiviral expression vector along with third-generation lentiviral packaging and pseudotyping plasmids (Dull et al., 1998) into the packaging cell line HEK293T. Briefly, 1.4 × 10^6^ cells were seeded in a 60mm tissue culture dish and grown to 80% confluence. 3µg of expression plasmid was co-transfected with lentiviral packaging and pseudotyping plasmids (2.25µg each of pMDLg/pRRE and pRSV-REV and 1.5µg of pMD2.G), using Lipofectamine 2000 (Invitrogen, Mulgrave, Vic, Australia) according to the manufacturer’s protocol. Cell culture medium was replaced after 24hr. The viral supernatant was collected 48hr post transfection and filtered using a 0.45µm filter. The filtered lentiviral supernatant was concentrated 20-fold by using Amicon Ultra-4 filter units (100 kDa NMWL) (Millipore, North Ryde, NSW, Australia).

### Lentiviral infection

4T1 cells were plated at a density of 1.0 × 10^5^ cells per well in 6-well tissue culture plates and culture medium was replaced after 24hr with medium containing 8µg/mL of polybrene (Sigma-Aldrich, Lismore, NSW, Australia). The cells were infected overnight with the concentrated virus at 1:5 dilution. Culture medium was changed 24hr post infection and cells were grown until reaching confluence. Cells transduced with both pSLIK-Venus-TmiR-Id1 and pSLIK-Neo-TmiR-Id3 were sorted on FACS using Venus as a marker followed by selection with neomycin at 400µg/mL for 5 days. Cells transduced with pSLIK-Neo-TmiR-EGFP were also selected with neomycin.

### Tumorsphere assay

Cells dissociated from modified 4T1 cells and p53-/- Id1/GFP, Id1C3-Tag tumors were put into tumorsphere assay as described previously (Nair et al., 2014a).

### Limiting dilution assay

Single-cell suspensions of FACS sorted Id1/GFP+ or unsorted viable tumor cells were prepared as described previously. Tumor cells were transplanted in appropriate numbers into the fourth mammary fat pad of 8- to 12-week-old FVB/N mice and aged till ethical end point. Extreme limiting dilution analysis software (Hu and Smyth, 2009) was used to calculate the TPF.

### *In vivo* and *ex vivo* imaging

The 4T1 cells were lentivirally modified with the pLV4311-IRES-Thy1.1 vector, a luciferase expressing vector (a kind gift from Dr Brian Rabinovich, The University of Texas M.D. Anderson Cancer Center, Houston, TX, USA). Animals were imaged twice weekly. Briefly, mice were first injected intraperitoneally with 200μL of 30% D-luciferin (Xenogen, Hopkinton, MA, USA) in PBS with calcium and magnesium (Life Technologies, Mulgrave, Vic, Australia) and imaged under anesthesia using the IVIS Imaging System 200 Biophotonic Imager (Xenogen, Alameda, CA, USA). Bioluminescent intensity was analysed and quantified using the Image Math feature in Living Image 3.1 software (Xenogen, Alameda, CA, USA). For *ex vivo* imaging, 200μL of 30% D-luciferin was injected into the mice just before autopsy. Tissues of interest were collected, placed into 6-well tissue culture plates in PBS, and imaged for 1–2min. At ethical endpoint, lungs were harvested and visually examined to detect the presence of metastases and later quantified based on 4T1 GFP fluorescence under a dissecting microscope.

### MTS proliferation assay

Cell viability assay (MTS assay) was carried out using the CellTiter 96 AQueous Cell Proliferation Assay (G5421; Promega, Alexandria, NSW, Australia) according to the manufacturer’s recommendations.

### Microarray and bioinformatics analysis

Total RNA from the samples were isolated using Qiagen RNeasy minikit (Qiagen, Doncaster, VIC, Australia. cDNA synthesis, probe labelling, hybridization, scanning and data processing were all conducted by the Ramaciotti Centre for Gene Function Analysis (The University of New South Wales). Gene expression profiling was performed using the Affymetrix GeneChip® Mouse Gene 2.0 ST Array. Normalization and probe-set summarization was performed using the robust multichip average method (Irizarry et al., 2003) implemented in the Affymetrix Power Tools apt-probeset-summarize software (version 1.15.0) (using the -a rma option). Differential expression between experimental groups was assessed using Limma (Smyth, 2004) via the limmaGP tool in GenePattern (Reich et al., 2006). Gene Set Enrichment Analysis (GSEA) (http://www.broadinstitute.org/gsea) (Subramanian et al., 2005) was performed using the GSEA Pre-ranked module on a ranked list of the limma moderated t-statistics, against gene-sets from v4.0 of the MSigDB (Subramanian et al., 2005) and custom gene-sets derived from the literature. Microarray data are freely available from GEO: GSE129790

### Next generation sequencing

3.5×10^4^ 4T1 K1 cells were seeded in 6-well plates in 4T1 media and treated with or without Doxorubicin (1 µg/mL) to induce Id1/3 knockdown. Cells were also transfected with non-targeting control siRNA (Dharmacon D-001810-10-05) or Robo1 siRNA (Dharmacon M-046944-01-0010). Cells were harvested after 48 hours and total RNA was extracted using the automated QiaSymphony magnetic bead extraction system. The Illumina TruSeq Stranded mRNA Library Prep Kit was used to generate libraries with 1 μg of input RNA following the manufacturer’s instructions. cDNA libraries were sequenced on the NextSeq system (Illumina), with 75 bp paired-end reads. Quality control was checked using FastQC (bioinformatics.babraham.ac.uk/projects/fastqc). Reads were then aligned to the mouse reference genome Mm10 using STAR ultrafast universal RNA-Seq aligner (Dobin et al., 2013). Gene feature counting was performed with RSEM (Li and Dewey, 2011). Replicate 3 from the Id1 KD group showed no KD of Id1 by qPCR and was therefore removed prior to down-stream differential expression analysis. Transcripts with expression counts of 0 across all samples were removed and then normalised using TMM (Robinson and Oshlack, 2010). The normalized counts were then log transformed using voom (Ritchie et al., 2015) and differential expression was performed with limma (Smyth, 2004). Differentially expressed genes were visualized and explored using Degust (http://degust.erc.monash.edu/). Genes with false discovery rate (FDR)<0.05 were considered significantly differentially expressed. For GSEA analysis, genes were ranked based on the limma moderated t-statistic and this was used as input for the GSEA desktop application (Subramanian et al., 2005). RNA sequencing data are freely available from GEO: GSE129858.

Microarray (GSE129790) and RNA-Seq (GSE129858) datasets are available in SuperSeries GSE129859.

### Statistical analysis

Statistical analyses were performed using GraphPad Prism 6. All *in vitro* experiments were done in 3 biological replicates each with 2 or more technical replicates. 5-10 mice were used per condition for the *in vivo* experiments. Data represented are means ± standard deviation. Statistical tests used are Unpaired student t-test and two-way-ANOVA. p-values <0.05 were considered statistically significant with \**p*< 0.05, \*\**p*< 0.01, *** *p*< 0.001, **** *p*< 0.0001.

## Results

### Id marks a subset of cells with stem-like properties in TNBC models

We investigated the role of Id in the context of CSC biology in the TNBC molecular subtype. Immunohistochemistry (IHC) analysis revealed that ID1 is expressed by a small minority of cells (range 0.5-6% of total cancer cells) in ∼50 % of ER-negative disease, namely TNBC and Her2+ tumors (Supplementary Figure S1A, B). No significant difference in the distribution of ID3 expression was observed across different subtypes (data not shown).

To test the hypothesis that Id1^+^ cells have a unique malignant phenotype, we developed two murine models of TNBC that permit the prospective isolation of Id1^+^ cells for functional assays. In the first, we used the p53-/- TNBC tumor model where IHC analysis revealed that ∼ 5% of neoplastic cells expressed Id1, consistent with the observation in the clinical samples, while Id3 marked a majority of the tumor cells in this model (Figure 1A).

**Figure 1.**
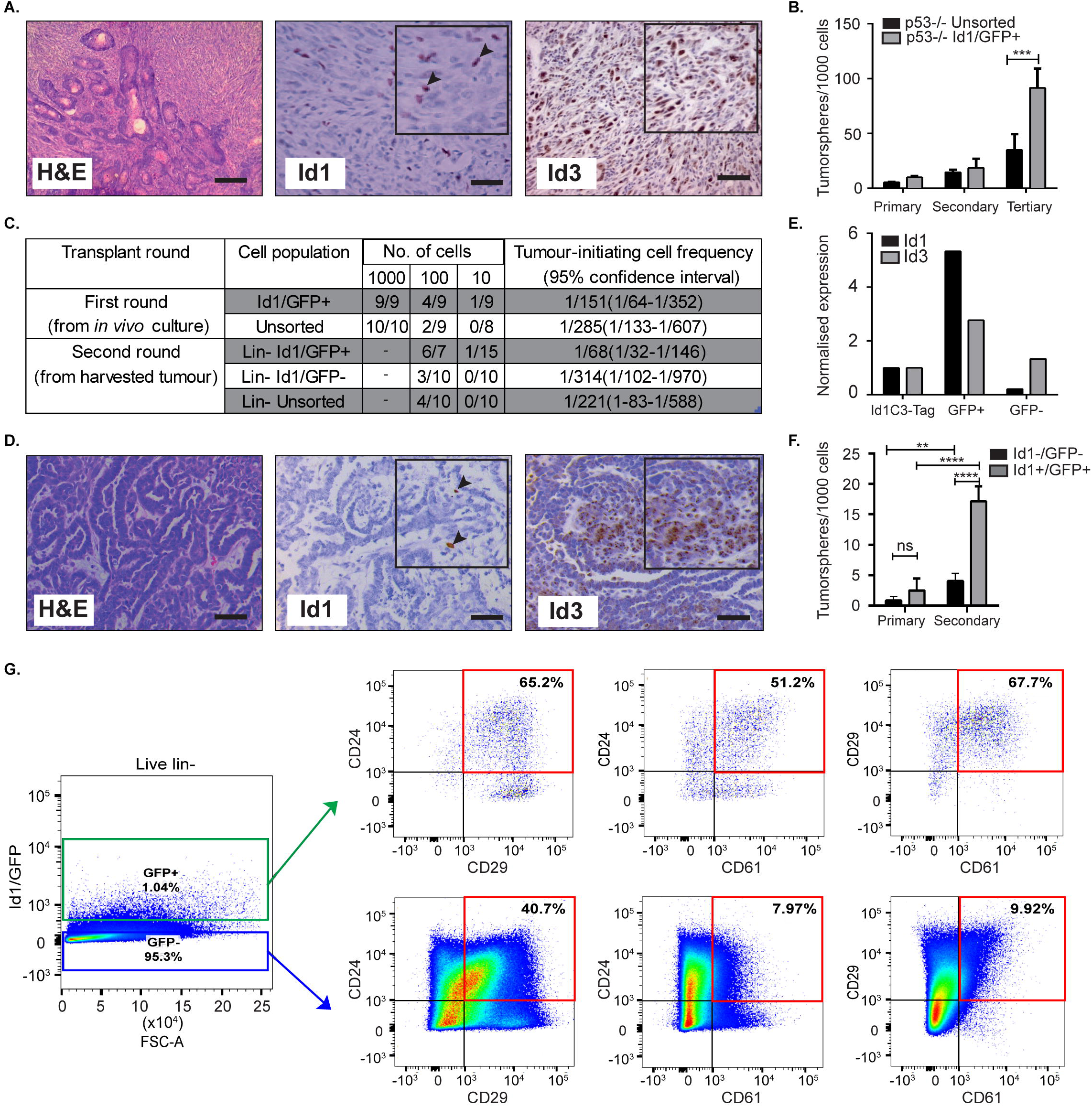
Id1 marks tumor cells with high self-renewal in murine models of TNBC. **(A)** Representative IHC images of Id1 and Id3 expression in p53-/- tumor model. Black arrows in the inset indicate Id1+ cells. Scale bars = 50μm. **(B)** p53-/- tumor cells were transfected with the Id1/GFP reporter and subsequently sorted for GFP expression. The self-renewal capacity of Id1/GFP^+^ p53-/- cells was significantly higher than unsorted Id1/GFP p53-/- cells upon passage to tertiary tumorspheres. Data are means ± SD (n=3). (*** *p*< 0.001; Two-way ANOVA). **(C)** Id1 expressing cells were sorted from the p53-/- Id1/GFP tumor model and transplanted into recipient mice by limiting dilution assay. Based on limiting dilution calculations (ELDA), the Id1^+^ cells demonstrated a 4.6-fold enrichment in tumor initiating capacity (TIC) when compared to the Id1^-^ cells in serial passage. p values for p53-/- Id1GFP+ versus Unsorted Round 1-0.2920, p53-/- Id1GFP+ versus Id1GFP-Round 2-0.0221**. (D)** Representative IHC images of the Id1C3-Tag model, confirming its suitability as a model system. Black arrows in the inset indicate Id1+ cells. Expression of Id1 was less than 5% as determined by IHC. Bars = 50μm. **(E)** Tumor cells from the Id1C3-Tag tumor model were FACS sorted based on their GFP expression. qRT-PCR analyses on the sorted GFP^+^ and GFP^-^ cell populations showed a significant increase (more than 5-fold) for Id1 expression in the GFP^+^ cells compared to cells lacking GFP expression. **(F)** *In vitro* self-renewal capacity of GFP^+^ cells was measured using the tumorsphere assay. The secondary sphere forming capacity of Id1^+^ tumor cells from the Id1C3-Tag model was significantly enriched in comparison to the Id1^-^tumor cells. Data are means ± SD (n=3). (\*\**p*< 0.01, **** *p*< 0.0001; Two-way ANOVA). **(G)** Representative FACS scatterplot and histograms from Id1C3-Tag tumors showing the expression of the CSC markers CD24, CD29 and CD61 in the Id1^-^/GFP^-^ and Id1^+^/GFP^+^ cancer cells. Putative CSC populations are highlighted within the red box.

To create a genetic reporter cell line, p53-/- mammary tumor cells were transduced with a lentiviral GFP reporter construct under the control of the Id1 promoter (Id1/GFP), as described previously (Mellick et al., 2010) (Supplementary Figure S1C). FACS sorting for GFP expression followed by immunoblotting confirmed the ability of the Id1/GFP construct to prospectively enrich for Id1+ cells from this model (Supplementary Figure S1D). We next sought to understand if Id1 marked cells with high self-renewal capacity in this model using tumorsphere assays, a well-established surrogate for cells with high self-renewal capacity (Lee et al., 2016a; Pastrana et al., 2011). We observed an increase in the self-renewal capacity of Id1/GFP^+^ cells when compared to the unsorted cell population in the p53-/- model (Figure 1B).

To establish the *in vivo* relevance of the increased self-renewal capacity of the Id1/GFP^+^ tumor cells observed *in vitro*, we determined the tumor initiating capacity (TIC) of the Id1/GFP^+^ cells using the limiting dilution assay (Nair et al., 2014a). Id1/GFP^+^ cells (1/68) showed more than a 4.6-fold increase in tumor initiating cell frequency over Id1/GFP^-^ cells (1/314) after serial passage (Figure 1C).

We used the Id1C3-Tag tumor model as a second murine model to assess the phenotype of Id1^+^ cells. In the C3-Tag tumor model, the expression of SV40-large T antigen in the mammary epithelium under the control of the C3 promoter leads to the development of TNBC in mice (Green et al., 2000; Pfefferle et al., 2013). These tumors (C3-Tag) closely model the TNBC subtype as assessed by gene expression profiling (Pfefferle et al., 2013). To generate a genetic reporter of Id1 promoter activity in TNBC, the C3-Tag model was crossed to a genetic reporter mouse model in which GFP is knocked into the intron 1 of the Id1 gene (Perry et al., 2007). The resulting Id1GFPC3-Tag mice (called Id1C3-Tag model) developed mammary tumors with similar kinetics as the parental C3-Tag mice and have a classical basal phenotype characterized by CK14^+^/CK8^-^ phenotype (Supplementary Figure S1E). 5% and 60% of cells in the Id1C3-Tag tumor were stained positive for Id1 and Id3 expression, respectively, as observed by IHC (Figure 1D). We were able to isolate Id1^+^ tumor cells with a high degree of purity by FACS based on GFP expression followed by q-RT PCR (Figure 1E). The sorted cells were put into primary tumorsphere assay and the spheres were serially passaged to secondary and tertiary spheres which robustly selects for self-renewing cell populations. Similar to the p53-/- Id1/GFP model, Id1^+^/GFP^+^ cells from the Id1C3-Tag model were enriched for sphere-forming capacity (Figure 1F).

Using the Id1C3-Tag model, we also looked at the association of Id1/GFP expression with the expression of established CSC markers CD29, CD24 and CD61. CD29^+^ /CD24^+^ status was previously reported to mark the tumorigenic subpopulation of cells in murine mammary tumors (Herschkowitz et al., 2012; Zhang et al., 2008). The Id1^+^/GFP^+^ cells in the Id1C3-Tag model are predominantly of the CD29^+^ /CD24^+^ phenotype (Figure 1G), with a 1.6-fold higher proportion of cells expressing both CD29 and CD24 compared to the Id1^-^/GFP^-^ cells which comprise the bulk of the tumor. Interestingly, Id1^+^/GFP^+^ cells are also highly enriched for CD24^+^/CD61^+^ expression (more than 6-fold increase in Id1^+^/GFP^+^ cells), which was also reported to mark a murine breast CSC population (Vaillant et al., 2008) (Figure 1G).

We found no correlation between Id1 expression (as indicated by GFP) and the CD29^+^ /CD24^+^ phenotype in the first transplantation round (T1) using the p53-/- model, as the percentage of CD29^+^ /CD24^+^ cells was similar across each gating group (Supplementary Figure S1F). Interestingly, the Id1^+^ cells, which are the putative cells that give rise to the increased TIC as shown in Figure 1C, showed 10 times less CD24^+^/CD29^+^ cells in the second transplantation round (T2) (Vaillant et al., 2008). The ability of the markers like CD24, CD29 and CD61 to identify the CSC population is clearly model-dependent. In addition to CD29 and CD24, the percentage of GFP^+^ cells were also analysed and a higher percentage of GFP^+^ cells was found in the second transplantation round of the p53-/- tumor compared to the first round tumor result (Supplementary Figure S1G), consistent with the increase in TICs reported in Figure 1C.

### Id requirement for self-renewal *in vitro* and metastatic competency *in vivo*

We next assessed the requirement for Id1 and Id3 in maintaining the CSC phenotypes. Numerous studies have shown that there exists a functional redundancy between Id1 and Id3, so studies typically require depletion of both the factors to reveal a phenotype (Konrad et al., 2017). Unfortunately we could not generate Id1 and Id3 double out knock mice for the C3Tag and Id1/3 expressing reporter in the p53-/- tumor models due to technical reasons. Hence we decided to look at the role of both Id1 and Id3 in the context of a knock down model. We used the transplantable syngeneic 4T1 TNBC model, which has a high propensity to spontaneously metastasize to distant sites (including bone, lung, brain and liver), mimicking the aggressiveness of human breast cancers (Aslakson and Miller, 1992; Eckhardt et al., 2005; Lelekakis et al., 1999; Pulaski and Ostrand-Rosenberg, 1998; Tao et al., 2008; Yoneda et al., 2000). IHC analysis showed that 15% of 4T1 tumor cells express high levels of Id1, and 35% have intermediate levels of Id1 expression, whereas the expression of Id3 was found in most of the cells (Figure 2A).

**Figure 2.**
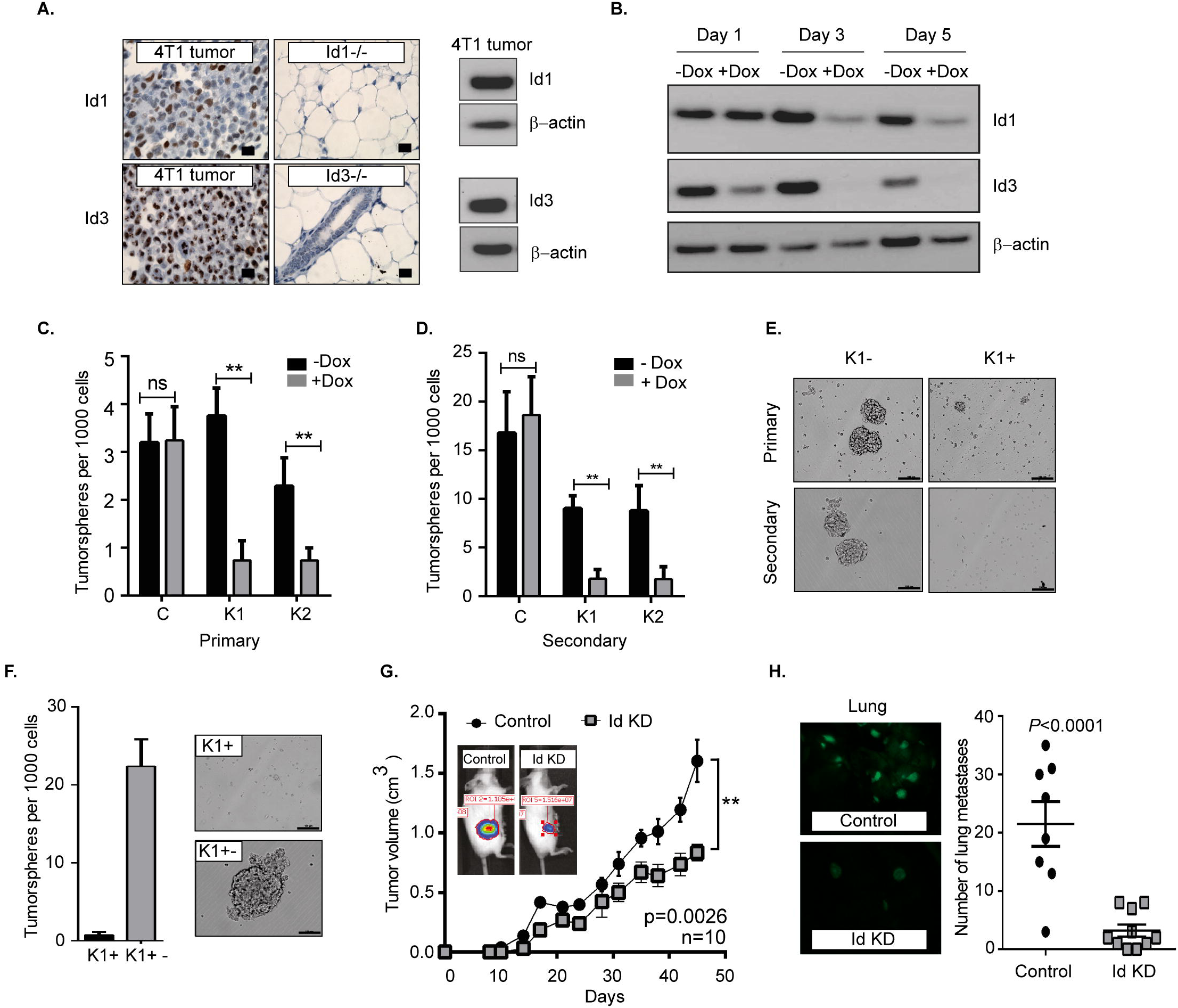
Depletion of Id1 and Id3 leads to a reduced self-renewal capacity *in vitro* and metastatic potential *in vivo*. **(A)** Endogenous levels of Id1 and Id3 expression in 4T1primary mammary tumors were determined. 4T1 were cells stained for Id1 and Id3 expression (brown) and counterstained with haematoxylin. Mammary gland tissue from Id1 and Id3 null (Id1-/- and Id3-/-) mice served as negative controls. Scale bars = 50 μm. Western blot analysis of protein lysate from 4T1 tumor cells served as positive controls for Id1 and Id3 expression**. (B)** Kinetics of conditional Id knockdown in 4T1 cells. Representative Western blot analysis of Id protein levels in pSLIK K1 cells over time. Cells were cultured in the presence of 1 μg/ml of Doxycycline (Dox) for 1, 3 and 5 days. β-actin was used as loading control. **(C)** 4T1 Control, pSLIK K1 and K2 clones were assayed for their tumorsphere forming potential. Dox was added into the culture medium at day 0. Number of primary tumorspheres formed was quantified by visual examination on day 7. Id knockdown leads to a decrease in tumorsphere-forming ability of K1 and K2 cell lines. Data are means ± SD (n=3). (\*\**p*< 0.01; Two-way ANOVA). **(D)** Primary tumorspheres were passaged and the number of secondary tumorspheres was quantified on day 14. Knockdown of Id significantly reduces the ability of the K1 and K2 cells to form secondary tumorspheres in the suspension culture. Data are means ± SD (n=3). (\*\**p*< 0.01; Two-way ANOVA). **(E)** Representative images of primary and secondary tumorsphere formation for the clone K1 ±Dox. **(F)** Quantification and representative images of primary tumorsphere treated with Dox (K1+) passaged to secondary spheres in Dox free conditions (K1+-) allowing re expression of Id and restoration of self-renewal capacity. **(G)** Knockdown of Id significantly delays tumor growth in the 4T1 syngeneic model. (*n* = 10 mice; **p value<0.01, Student’s *t*-test). **(H)** Id knockdown suppresses spontaneous lung metastasis. Tumors depleted of Id expression generated fewer spontaneous lung macrometastatic lesions compared to the control despite growing in the host for a longer time. Inset shows representative images of lungs bearing the control (K1 - Dox) and Id KD (K1 + Dox) lung metastases at ethical end point. Control; n = 8 mice, Id KD; n=10 mice. Scale bar = 50 um.

We next used an inducible lentiviral shRNA system (Shin et al., 2006) that permits reversible knock down of Id1 and Id3 in response to doxycycline (Dox) treatment in 4T1 cells. Two clonal 4T1 cell lines, K1 and K2 were chosen along with a control line (C), based on the efficiency of Id knock down (Figure 2B, Supplementary Figure S2A). Id depletion resulted in a significant decrease in cell proliferation and migration *in vitro* when compared to the control (Supplementary Figure S2 B, C, D).

We next interrogated the effect of Id depletion on the self-renewal capacity of the C, K1 and K2 cell lines. Dox-dependent shRNA induction significantly reduced the ability of the K1 and K2 cells to form primary tumorspheres in the suspension culture (Figure 2C). This effect was not observed in the control cell line (C; Figure 2C). A significant further decrease in self-renewal capacity of K1 and K2 lines was observed when primary tumorspheres were passaged to the secondary stage (Figure 2D, E). The Id depleted tumorspheres were also markedly smaller in size compared to controls (Figure 2E, Supplementary Figure S2E).

To assess if the self-renewal phenotype controlled by Id is reversible, we firstly passaged primary tumorspheres [previously treated with Dox (K+)] to secondary tumorspheres. The secondary tumorspheres were then cultured in the presence or absence of Dox, to maintain the Id knockdown status or to allow the re-expression of Id, respectively (Supplementary Figure S2F, G). The secondary tumorspheres cultured without Dox (K1+-) re-established their self-renewal capacity as evidenced by the ability to form new tumorspheres (Figure 2F; Supplementary Figure S2H, I), suggesting that Id depletion does not lead to a permanent loss of self-renewal capacity.

To determine whether Id1 and Id3 are required for primary tumor and metastatic growth *in vivo*, K1 cells were orthotopically transplanted into the mammary fat pad of BALB/c mice. Dox-mediated knockdown of Id resulted in modest inhibition of primary tumor growth, with control tumors growing faster and reaching the ethical endpoint earlier than the Id knockdown group (Figure 2G). More significantly, mice transplanted with Id depleted K1 cells presented far fewer lung metastatic lesions compared to the control despite growing in the host for a longer time (p<0.0001; Figure 2H).

To assess the role for Id in metastatic progression *in vivo*, we examined Id expression in lung metastasis compared to primary tumors in mice injected with K1 cells. An increase in the expression of Id1 was observed in the lung metastasis in all the samples, while no significant enrichment of Id3 expression was observed (Supplementary Figure S3). This suggests that Id1 promotes lung metastatic dissemination in TNBC.

### Identification of genes and pathways regulated by Id

The canonical role for Id proteins is to regulate gene expression through association with transcription factors, yet a comprehensive analysis of Id transcriptional targets in cancer has not been reported. We performed gene expression profiling of Control (C) and Id depleted K1 cells. The gene expression profiles of four independent replicates (R1, R2, R3 and R4 ± doxycycline treatment) were compared by microarray analysis (Supplementary Figure S4A). 6081 differentially expressed genes were identified (Q<0.05), with 3310 up-regulated and 2771 down-regulated genes in Id KD cells (Supplementary Table S1) shows the top 25 differentially regulated genes). Network and pathway enrichment analysis was conducted using the MetaCore^™^ software. 4301 significant network objects were identified for the Id knockdown microarray data (adjusted p-value of ≤0.05). The top pathways affected by Id knockdown were mostly associated with the cell cycle (Figure 3A, B) consistent with the loss of proliferative phenotype described previously (Supplementary Figure S2B, C). Similar results were obtained using Gene Set Enrichment Analysis (GSEA) with significant down regulation of proliferative signatures (CELL_CYCLE_PROCESS) and mitosis (M_PHASE) (Supplementary Table S2). Genes such as CCNA2, CHEK1 and PLK1 in these gene sets are down-regulated by Id knockdown. This is consistent with our results (Supplementary Figure S2 B, C) showing Id proteins are necessary for proliferation of 4T1 cells, as well as previous studies which reported a role of Id in controlling cell cycle progression and proliferation pathways (Nair et al., 2014b; O’Brien et al., 2012). Enrichment for genes involved in several oncogenic pathways such as Mek, Vegf, Myc and Bmi1 signalling have also been highlighted (Supplementary Table S3). In order to identify whether Id specifically regulate genes controlling breast cancer metastasis, GSEA analysis was performed with a collection of custom “metastasis gene sets”. This collection (Table 1) consists of several metastatic signatures from the C2 collection (MSigDB database; Supplementary Table S4), combined with a list of custom gene sets described in major studies (Aceto et al., 2014; Bos et al., 2009; Charafe-Jauffret et al., 2009; Dontu et al., 2003; Kang et al., 2003b; Liu et al., 2010; Minn et al., 2005a; Minn et al., 2005b; Padua et al., 2008; Tang et al., 2007) as shown in Figure 3C. Genes differentially expressed in this set included *Robo1* (Chang et al., 2012; Qin et al., 2015), *Il6* (Chang et al., 2013), *Fermt1* (Landemaine et al., 2008), *Foxc2* (Mani et al., 2007) and Mir30a (Zhang et al., 2014). Three putative Id targets *Robo1, Fermt1* and *Mir30a* were then validated using q-RT PCR (Figure 3D) and found to be differentially regulated in the K1 cell line upon Id KD.

**Table 1.** Gene expression signatures of breast cancer metastasis and breast cancer stem cells. This table showed a collection of gene sets which comprised several metastatic signatures that were picked from the C2 collection on the MSigDB database and several other signatures that were manually curated. GSEA analysis was carried out to identify whether any of the Id1/3 targets from the profiling experiment are enriched in these signatures.

| Study | Signature | Type | Available on GSEA MSigDB database? |
| --- | --- | --- | --- |
| Landemaine et al. A six-gene signature predicting breast cancer lung metastasis. Cancer Res. 2008 Aug 1;68(15):6092-9 | Lung metastasis signature of breast cancer | Metastatic tissue tropism | Yes |
| Bild et al. Oncogenic pathway signatures in human cancers as a guide to targeted therapies. *Nature* 2006, 439:353-357. | Expression profile of 4 individual genes -- Myc, E2F3, Ras, Src, β-catenin | Signalling pathway | Yes |
| van 't Veer et al. Gene expression profiling predicts clinical outcome of breast cancer. *Nature* 2002, 415:530-536. | Poor prognosis signature of breast cancer | Classifier that classifies patients as having good or poor prognosis | Yes |
| Wang et al. Gene-expression profiles to predict distant metastasis of lymph-node-negative primary breast cancer. *Lancet* 2005, 365:671-679. | Poor prognosis signature of breast cancer | Classifier | Yes |
| Ramaswamy et al. A molecular signature of metastasis in primary solid tumors. *Nat Genet* 2003, 33:49-54. | General metastasis | Classifier | Yes |
| Finak et al. Stromal gene expression predicts clinical outcome in breast cancer. *Nat Med* 2008, 14:518-527. | Breast tumor stromal gene expression signature | Classifier | Yes |
| Farmer et al. A stroma-related gene signature predicts resistance to neoadjuvant chemotherapy in breast cancer. *Nat Med* 2009, 15:68-74. | Stromal gene expression signature of breast tumor treated with chemotherapy | Classifier | Yes |
| Kang et al. A multigenic program mediating breast cancer metastasis to bone. *Cancer Cell* | Bone metastasis signature of breast | Metastatic tissue tropism | No |
| 2003, 3:537-549. | cancer |  |  |
| Minn et al. Genes that mediate breast cancer metastasis to lung. Nature 2005, 436:518-524 | Lung metastasis signature of breast cancer | Metastatic tissue tropism | No |
| Bos et al. Genes that mediate breast cancer metastasis to the brain. Nature 2009, 459:1005-1009. | Bone metastasis signature of breast cancer | Metastatic tissue tropism | No |
| Padua et al. TGFbeta primes breast tumors for lung metastasis seeding through angiopoietin-like 4. Cell 2008, 133:66-77. | TGF-b signature in lung metastasis of breast cancer | Signalling pathway | No |
| Aceto et al. Tyrosine phosphatase SHP2 promotes breast cancer progression and maintains tumor-initiating cells via activation of key transcription factors and a positive feedback signaling loop. Nat Med. 2012 Mar 4;18(4):529-37. | Shp2 signature in breast cancer metastasis | Signalling pathway | No |
| Minn et al. Distinct organ-specific metastatic potential of individual breast cancer cells and primary tumors. J Clin Invest. 2005 Jan;115(1):44-55. | Poor prognosis signature of breast cancer ; Breast cancer metastasis signature; Bone metastasis signature of breast cancer | Metastatic tissue tropism | No |
| Tang et al. Transforming growth factor-beta can suppress tumorigenesis through effects on the putative cancer stem or early progenitor cell and committed progeny in a breast cancer xenograft model. Cancer Res. 2007 Sep 15;67(18):8643-52. | TGF-b signature in lung metastasis of breast cancer | Signalling pathway | No |
| Liu et al. The prognostic role of a gene signature from tumorigenic breast-cancer cells. The New England journal of medicine. 2007. 356(3), 217-26. | Gene signatures of CD44+CD24-/low tumorigenic breast-cancer cell-lines and normal breast epithelium | Cancer stem cell | No |
| Charafe-Jauffret et al. Breast cancer cell lines contain functional cancer stem cells with | Breast cancer stem cell signature | Cancer stem cell | No |
| metastatic capacity and a distinct molecular signature. 2009. Cancer research, 69(4), 1302-13. |  |  |  |
| Dontu, et al. In vitro propagation and transcriptional profiling of human mammary stem/progenitor cells. 2003. Genes & development, 17(10), 1253-70. | Gene signature of human mammary stem and progenitor cells | Cancer stem cell/ Differentiation | No |

**Figure 3.**
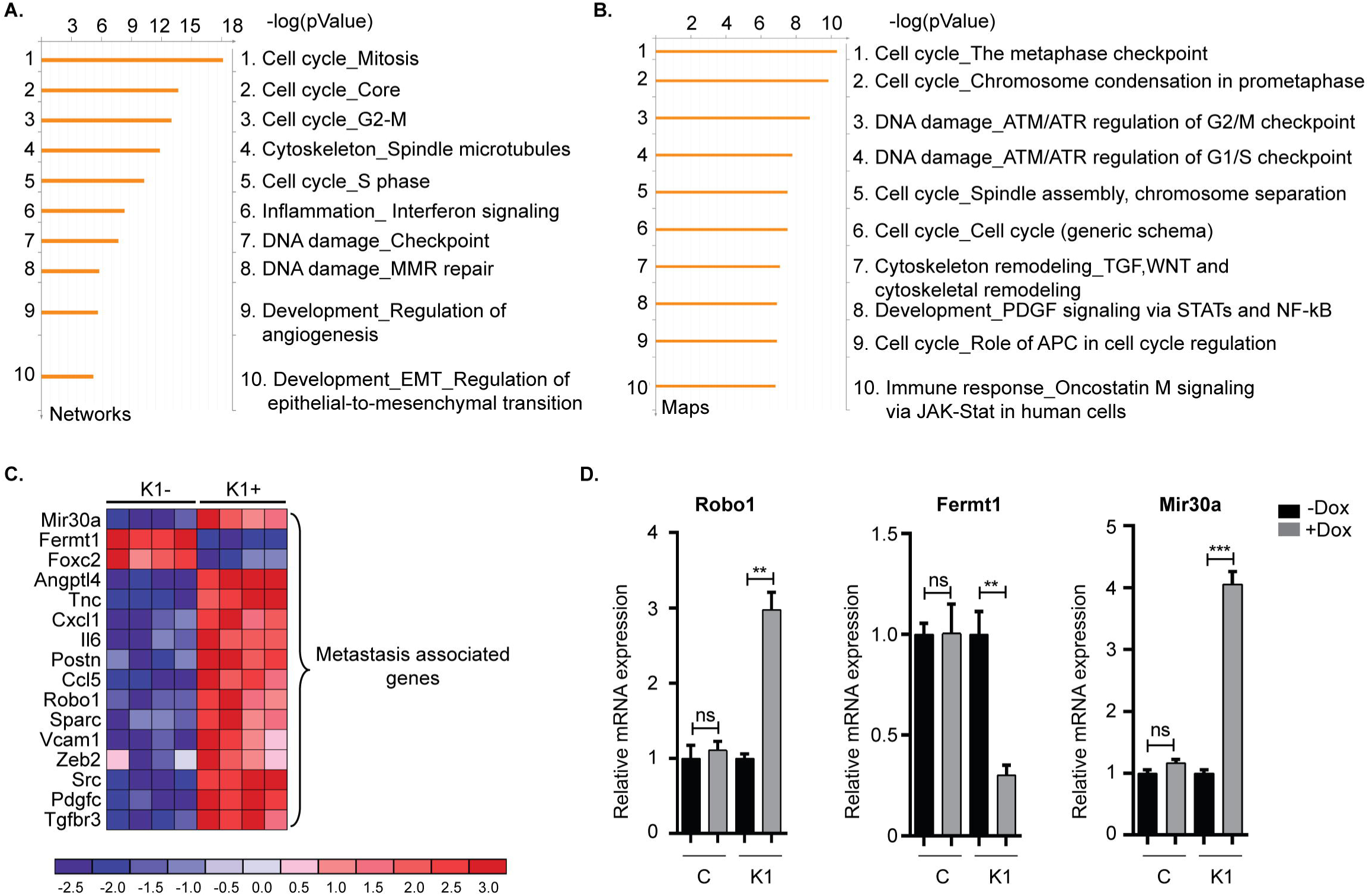
Gene expression analysis reveals targets of Id in TNBC. **(A, B)** To characterize the network of genes regulated by Id, functional annotation analyses were performed on the gene array data from the 4T1 TNBC model. The Id depletion model attempted to identify downstream targets of Id through a loss of function approach. The gene expression profile of four independent replicates of the K1 shId clone, with and without doxycycline treatment, was compared by microarray analysis. This resulted in a list of differentially expressed genes between control and Id depleted cells, which by further network and map analysis using Metacore demonstrated was largely driven by genes controlling cell cycle pathways. **(C)** Gene expression analysis identified metastasis-related genes that were differentially expressed in response to Id knockdown. To determine if genes that mediate metastasis were enriched in the Id signature, gene expression analysis was performed using a manually curated set of metastasis gene sets. Genes differentially expressed in response to Id knockdown as well as associated with pathways regulating metastasis were identified based on reports from the literature which included Robo1. **(D)** Validation of expression profiling results by quantitative real-time-PCR using the Taqman® probe based system. Relative mRNA expression of Robo1, Fermt1 and Mir30a, in the 4T1 pSLIK shId Clonal cell line (K1) and pSLIK control (C), as indicated. Data are means ± SD (n=3). (\*\**p*< 0.01, **** *p*< 0.0001; unpaired t-test).

### Id mediated inhibition of Robo1 controls the proliferative phenotype via activation of Myc transcription

Since Robo1 is known to have a tumor suppressor role in breast cancer biology (Chang et al., 2012; Shen et al., 2015), we next sought to determine if Robo1 has an epistatic interaction with Id loss of function using siRNA mediated knockdown of Robo1 followed by proliferation assays. Knockdown of *Robo1* ameliorated the requirement for Id and rescued approximately 55 % of the proliferative decrease induced by Id KD (Figure 4A).

To understand the mechanisms by which *Robo1* increases the proliferative potential of Id depleted cells *in vitro*, we performed RNA-Sequencing (RNA-Seq) experiments on K1 cells with dox-inducible Id KD and/or Robo1 depletion using siRNA. Four replicates per condition were generated and MDS plots presented in Supplementary Figure S4B showed that the replicates cluster together. Id KD alone in the K1 cells down regulated 4409 genes and up regulated 5236 genes (FDR<0.05), respectively. The majority of the differentially expressed genes determined by microarray were found by RNA-Seq analysis (Supplementary Figure S4C). Id depletion led to an increase in *Robo1* expression, as observed in the previous microarray experiment (Figure 3C, D; Figure 4 B).

**Figure 4.**
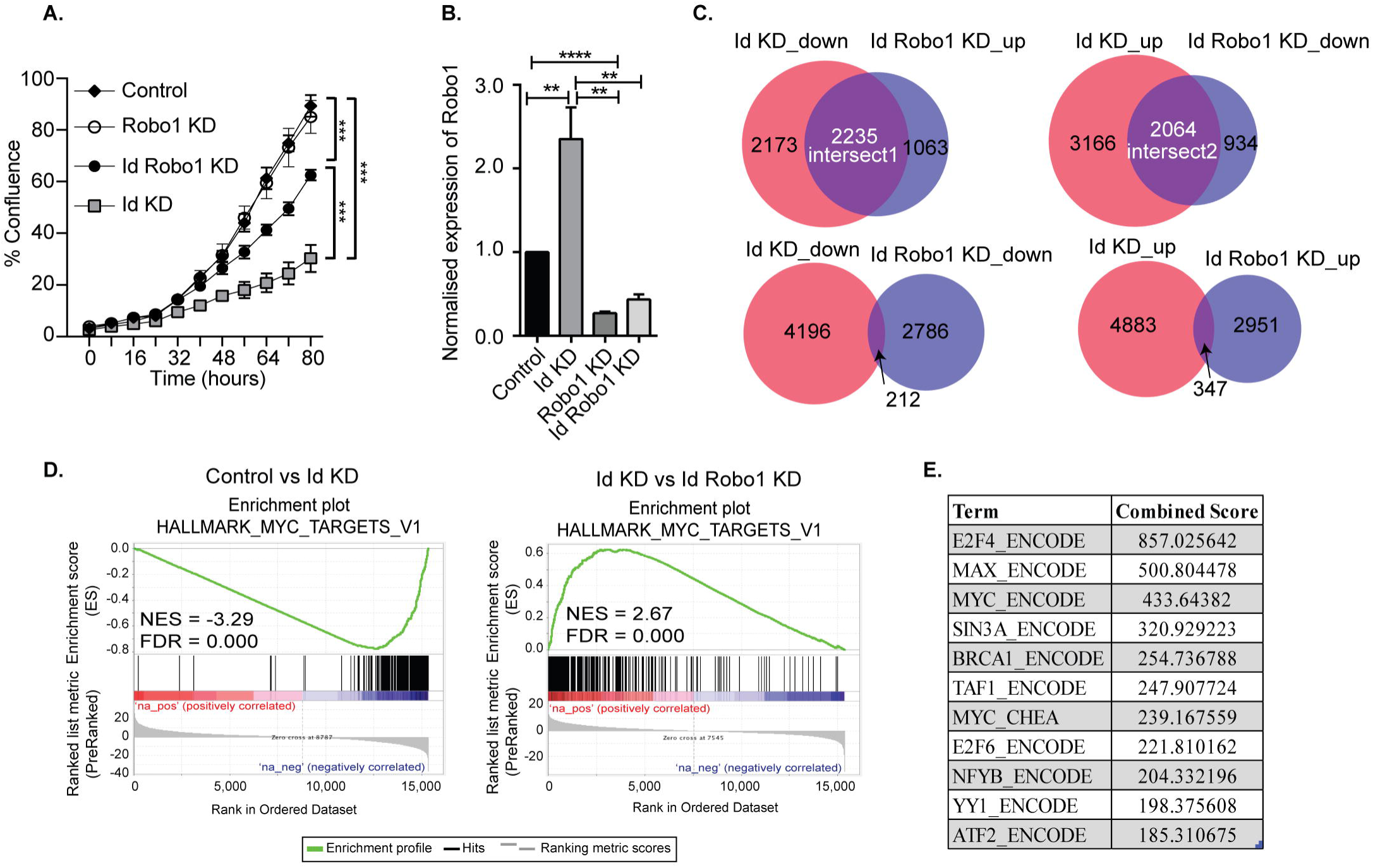
Identification of Myc signature activation by Id via negative regulation of Robo1. **(A)** Proliferation of K1 cells treated with non-targeting (NT) control siRNA or Robo1 siRNA in the absence or presence of Doxycycline to induce Id knockdown was measured by the IncuCyte^™^ (Essen Instruments) live-cell imaging system. Data shown as mean ±SD (n=3). (*** p< 0.001; Unpaired two-tailed t-test). **(B)** Robo1 expression in Control, Id KD, Robo1 KD and Id Robo1 KD cells was measured by quantitative PCR. Ct values were normalised to β actin and GAPDH housekeeping genes. Data shown as mean ± SEM (n=4). (** p< 0.01, **** p < 0.0001; Unpaired two-tailed t-test). **(C)** Transcriptional profiling was performed on Control, Id KD, Robo1 KD and Id Robo1 KD cells. Proportional Venn diagrams (BioVenn) were generated to visualise the overlapping genes between the different comparisons. **(D)** GSEA Enrichment plots of the hallmark Myc targets version 1 signature from MSigDB. NES = normalised enrichment score. **(E)** Consensus Transcription factor motif analysis using the Encyclopedia of DNA Elements (ENCODE) and ChIP enrichment analysis (ChEA) data sets determined using EnrichR. The combined score is a combination of the p-value and z-score.

Given that Id repressed Robo1 expression, we sought to determine Robo1 target genes in the absence of Id. Remarkably, under Id depletion conditions, *Robo1* KD restored expression of a large subset (∼45%) of Id target genes to basal levels (Figure 4C). In comparison, knockdown of Id or Robo1 regulated few targets in the same direction (e.g. both up or both down). This implies that a large proportion of Id targets may be regulated via suppression of Robo1. Genes whose expression was repressed by Id KD and rescued by concomitant Robo1 KD were termed ‘Intersect 1’ (Figure 4C, Table 2). Genes that were upregulated by Id KD and downregulated by Robo1 KD (in the absence of Id) were annotated ‘intersect 2’ (Figure 4C, Table 3). To investigate the function of these intersect group of genes, we performed GSEA analysis using the MSigDB hallmark gene set (Liberzon et al., 2015). The top signatures in Intersect 1 were involved in cell proliferation, with enrichment for G2M checkpoint, E2F and Myc targets as well as mTOR signalling (Table 2). Rank-based analysis revealed strong negative enrichment for the hallmark Myc targets signature upon Id knockdown alone, and strong positive enrichment upon Id and Robo1 knockdown (Figure 4D). This suggests that following Id KD, Robo1 is induced and exerts anti-proliferative effects via suppression of Myc and its target genes (Supplementary Figure S4D, E). Transcription factor motif analysis using EnrichR revealed that Myc and its binding partner Max, have a high combined score in the Intersect 1 gene list further implicating Myc as downstream effector of Robo1 and Id (Figure 4 E).

**Table 2.** GSEA on the Intersect 1 genes from Figure 4C against the MSigDB hallmark gene sets.

| **INTERSECT 1 GSEA** |  |  |  |  |  |  |
| --- | --- | --- | --- | --- | --- | --- |
| **Gene Set Name** | **# Genes in Gene Set (K)** | **Description** | **#Genes in Overlap (k)** | **k/K** | **p-value** | **FDR (q-value)** |
| HALLMARK_E2F_TARGETS | 200 | Genes encoding cell cycle related targets of E2F transcription factors. | 158 | 0.79 | 5.02E-178 | 2.51E-176 |
| HALLMARK_G2M_CHECKPOINT | 200 | Genes involved in the G2/M checkpoint, as in progression through the cell division cycle. | 116 | 0.58 | 2.32E-105 | 5.80E-104 |
| HALLMARK_MYC_TARGETS_V1 | 200 | A subgroup of genes regulated by MYC - version 1 (v1). | 113 | 0.565 | 7.79E-101 | 1.30E-99 |
| HALLMARK_OXIDATIVE_PHOSPHORYLATION | 200 | Genes encoding proteins involved in oxidative phosphorylation. | 96 | 0.48 | 1.04E-76 | 1.30E-75 |
| HALLMARK_MYC_TARGETS_V2 | 58 | A subgroup of genes regulated by MYC - version 2 (v2). | 42 | 0.7241 | 4.60E-45 | 4.60E-44 |
| HALLMARK_MTORC1_SIGNALING | 200 | Genes up-regulated through activation of mTORC1 complex. | 66 | 0.33 | 1.70E-40 | 1.42E-39 |
| HALLMARK_MITOTIC_SPINDLE | 200 | Genes important for mitotic spindle assembly. | 60 | 0.3 | 2.76E-34 | 1.97E-33 |
| HALLMARK_DNA_REPAIR | 150 | Genes involved in DNA repair. | 51 | 0.34 | 2.54E-32 | 1.59E-31 |
| HALLMARK_UNFOLDED_PROTEIN_RESPONSE | 113 | Genes up-regulated during unfolded protein response, a cellular stress response related to the endoplasmic reticulum. | 31 | 0.2743 | 3.62E-17 | 2.01E-16 |
| HALLMARK_FATTY_ACID_METABOLISM | 158 | Genes encoding proteins involved in metabolism of fatty acids. | 35 | 0.2215 | 5.10E-16 | 2.55E-15 |
| HALLMARK_ADIPOGENESIS | 200 | Genes up-regulated during adipocyte differentiation (adipogenesis). | 39 | 0.195 | 1.08E-15 | 4.89E-15 |
| HALLMARK_CHOLESTEROL_HOMEOSTASIS | 74 | Genes involved in cholesterol homeostasis. | 24 | 0.3243 | 1.95E-15 | 8.11E-15 |
| HALLMARK_ESTROGEN_RESPONSE_LATE | 200 | Genes defining late response to estrogen. | 35 | 0.175 | 8.70E-13 | 3.11E-12 |
| HALLMARK_GLYCOLYSIS | 200 | Genes encoding proteins involved in glycolysis and gluconeogenesis. | 35 | 0.175 | 8.70E-13 | 3.11E-12 |
| HALLMARK_UV_RESPONSE_UP | 158 | Genes up-regulated in response to ultraviolet (UV) radiation. | 28 | 0.1772 | 1.18E-10 | 3.95E-10 |
| HALLMARK_SPERMATOGENESIS | 135 | Genes up-regulated during production of male gametes (sperm), as in spermatogenesis. | 25 | 0.1852 | 4.39E-10 | 1.37E-09 |
| HALLMARK_ANDROGEN_RESPONSE | 101 | Genes defining response to androgens. | 20 | 0.198 | 7.15E-09 | 2.10E-08 |
| HALLMARK<br>_ESTROGEN<br>_RESPONSE<br>_EARLY | 200 | Genes defining early response to estrogen. | 28 | 0.14 | 2.76E-08 | 7.68E-08 |
| HALLMARK<br>_IL2_STAT5<br>_SIGNALIN<br>G | 200 | Genes up-regulated by STAT5 in response to IL2 stimulation. | 25 | 0.125 | 1.31E-06 | 3.27E-06 |
| HALLMARK<br>_KRAS_SIG<br>NALING_UP | 200 | Genes up-regulated by KRAS activation. | 25 | 0.125 | 1.31E-06 | 3.27E-06 |

**Table 3.** GSEA on the Intersect 2 genes from Figure 4C against the MSigDB hallmark gene sets.

| **INTERSECT**<br>**2 GSEA** |  |  |  |  |  |  |
| --- | --- | --- | --- | --- | --- | --- |
| **Gene Set**<br>**Name** | **#**<br>**Genes**<br>**in**<br>**Gene**<br>**Set**<br>**(K)** | **Description** | **# Genes**<br>**in**<br>**Overlap**<br>**(k)** | **k/K** | **p-value** | **FDR (q-**<br>**value)** |
| HALLMARK<br>_INTERFER<br>ON_GAMMA<br>_RESPONSE | 200 | Genes up-regulated in response to IFNG [GeneID=3458]. | 60 | 0.3 | 3.53E-38 | 1.76E-36 |
| HALLMARK<br>_INTERFER<br>ON_ALPHA_<br>RESPONSE | 97 | Genes up-regulated in response to alpha interferon proteins. | 43 | 0.4433 | 4.26E-36 | 1.06E-34 |
| HALLMARK<br>_HYPOXIA | 200 | Genes up-regulated in response to low oxygen levels (hypoxia). | 39 | 0.195 | 5.11E-18 | 8.51E-17 |
| HALLMARK<br>_P53_PATH<br>WAY | 200 | Genes involved in p53 pathways and networks. | 33 | 0.165 | 2.66E-13 | 3.33E-12 |
| HALLMARK<br>_APOPTOSIS | 161 | Genes mediating programmed cell death (apoptosis) by activation of caspases. | 28 | 0.1739 | 4.49E-12 | 4.49E-11 |
| HALLMARK<br>_ESTROGEN<br>_RESPONSE<br>_EARLY | 200 | Genes defining early response to estrogen. | 31 | 0.155 | 7.51E-12 | 4.70E-11 |
| HALLMARK<br>_HEME_ME<br>TABOLISM | 200 | Genes involved in metabolism of heme (a cofactor consisting of iron and porphyrin) and erythroblast differentiation. | 31 | 0.155 | 7.51E-12 | 4.70E-11 |
| HALLMARK<br>_MYOGENE<br>SIS | 200 | Genes involved in development of skeletal muscle (myogenesis). | 31 | 0.155 | 7.51E-12 | 4.70E-11 |
| HALLMARK<br>_PROTEIN_S<br>ECRETION | 96 | Genes involved in protein secretion pathway. | 21 | 0.2188 | 2.31E-11 | 1.28E-10 |
| HALLMARK<br>_EPITHELIA<br>L_MESENC<br>HIMAL_TRA<br>NSITION | 200 | Genes defining epithelial-mesenchymal transition, as in wound healing, fibrosis and metastasis. | 30 | 0.15 | 3.77E-11 | 1.89E-10 |
| HALLMARK<br>_ESTROGEN<br>_RESPONSE<br>_LATE | 200 | Genes defining late response to estrogen. | 29 | 0.145 | 1.82E-10 | 8.29E-10 |
| HALLMARK<br>_APICAL_JU<br>NCTION | 200 | Genes encoding components of apical junction complex. | 28 | 0.14 | 8.47E-10 | 3.53E-09 |
| HALLMARK<br>_IL2_STAT5<br>_SIGNALIN<br>G | 200 | Genes up-regulated by STAT5 in response to IL2 stimulation. | 26 | 0.13 | 1.62E-08 | 5.77E-08 |
| HALLMARK<br>_KRAS_SIG<br>NALING_DN | 200 | Genes down-regulated by KRAS activation. | 26 | 0.13 | 1.62E-08 | 5.77E-08 |
| HALLMARK<br>_ALLOGRAF<br>T_REJECTIO<br>N | 200 | Genes up-regulated during transplant rejection. | 25 | 0.125 | 6.63E-08 | 2.21E-07 |
| HALLMARK<br>_UNFOLDED<br>_PROTEIN_R | 113 | Genes up-regulated during unfolded protein response, a cellular stress | 18 | 0.1593 | 1.16E-07 | 3.62E-07 |
| ESPONSE |  | response related to the endoplasmic reticulum. |  |  |  |  |
| HALLMARK_ADIPOGENESIS | 200 | Genes up-regulated during adipocyte differentiation (adipogenesis). | 24 | 0.12 | 2.60E-07 | 6.83E-07 |
| HALLMARK_TNFA_SIGNALING_VIA_NFKB | 200 | Genes regulated by NF-kB in response to TNF [GeneID=7124]. | 24 | 0.12 | 2.60E-07 | 6.83E-07 |
| HALLMARK_XENOBIOTIC_METABOLISM | 200 | Genes encoding proteins involved in processing of drugs and other xenobiotics. | 24 | 0.12 | 2.60E-07 | 6.83E-07 |
| HALLMARK_IL6_JAK_STAT3_SIGNALING | 87 | Genes up-regulated by IL6 [GeneID=3569] via STAT3 [GeneID=6774], e.g., during acute phase response. | 15 | 0.1724 | 4.51E-07 | 1.13E-06 |

We were interested in investigating the possibility that Robo1 may exert its negative effects on the Myc pathway via regulation of Myc co-factors, which can potently enhance or suppress Myc transcriptional activity (Gao et al., 2016). In order to test this hypothesis, we looked at known Myc co-factors from the literature in our RNA-Seq data to determine if they were differentially expressed in the Id1 and Robo1 KD conditions. As seen in Supplementary Table S5, we included negative (red) and positive (green) cofactors in the analysis. Scrutiny of this list suggests that there are numerous negative co-factors (7/10) being induced and activators being repressed (13/24) by Robo1. For example, putative activation of the gene Rlim which is an E3 ubiquitin ligase that suppresses the transcriptional activity of MYC (Gao et al., 2016).

In summary, we have demonstrated that Id depletion leads to a loss in the proliferative and self-renewal cancer stem cell phenotypes associated with TNBC. Id1 acts by negatively regulating Robo1 which in turn finally leads to the downstream activation of a Myc transcriptional program (Figure 5).

**Figure 5:**
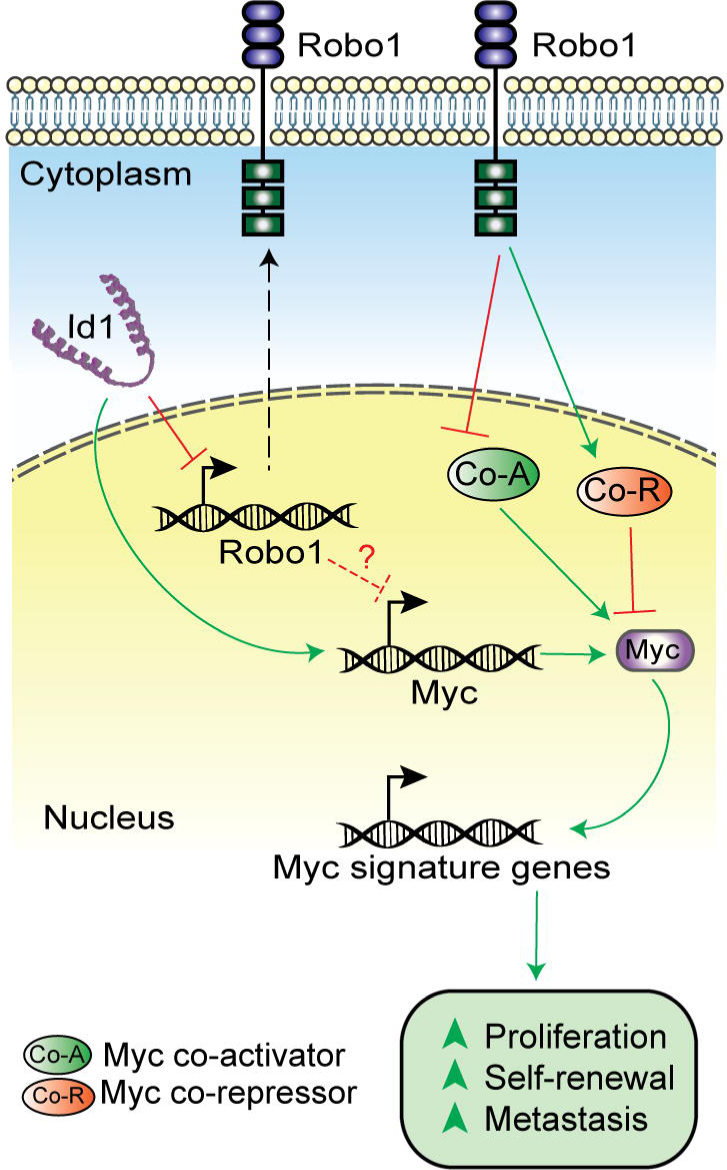
Model showing the mechanism of Id-Robo1 action in cancer cells. The proposed model for the regulation of Myc by Id and Robo1. Co-A indicates representative Myc activator and Co-R indicates representative Myc repressor.

## Discussion

There is increasing evidence that all cells within a tumor are not equal with some cells having the plasticity to adapt and subvert cellular and molecular mechanisms to be more tumorigenic than others. In this study, we demonstrate that Id1 and its closely related family member Id3 are important for the CSC phenotype in the TNBC subtype. Using four independent models of Id expression and depletion, we demonstrate that the properties of proliferation and self-renewal are regulated by Id proteins.

Transcription factors like the Id family of proteins can affect a number of key molecular pathways, allowing switching of phenotypes in response to local cues such as transforming growth factor-β (TGF-β) (Kang et al., 2003a; Stankic et al., 2013), receptor tyrosine kinase signalling (Tam et al., 2008), and steroid hormones (Lin et al., 2000) and therefore are able to transduce a multitude of cues into competency for proliferation and self-renewal. The CSC phenotype as marked by Id is plastic, fitting with the latest evidence that CSC are not necessarily hierarchically organised, but rather represent a transient inducible state dependent on the local microenvironment.

We report the first comprehensive analysis of Id transcriptional targets. We go on to identify a novel epistatic relationship with Robo1, with Robo1 loss sufficient to remove the necessity for Id in proliferation, suggesting that suppression of Robo1 is an important function for Id in this setting. Robo1 is a receptor for SLIT1 and SLIT2 that mediates cellular responses to molecular guidance cues in cellular migration (Huang et al., 2015). Previous work with mammary stem cells showed that the extracellular SLIT2 signals via ROBO1 to regulate the asymmetric self-renewal of basal stem cells through the transcription factor Snail during mammary gland development (Ballard et al., 2015). Our finding may have significant implications for tumor biology because SLIT/ROBO signalling is altered in about 40% of basal breast tumors (Ballard et al., 2015). Our work implicates a novel role for SLIT-ROBO signalling in CSC and shows a new mechanism by which Id proteins control the self-renewal phenotype by suppressing the Robo1 tumor suppressor role in TNBC.

The significant decrease in the Myc levels on Id knockdown suggest an Id/Robo1/Myc axis in TNBC (Supplementary Figure S4D, E). While the proposed model for regulation of Myc is not yet clear, we propose two possible modes of regulation of Myc: (1) Robo independent suppression of Myc expression and (2) Robo dependent regulation of Myc activity. Though the mechanism still needs to be elaborated, we hypothesise that in the absence of Id, Robo1 inhibits Myc activity via activation of Myc inhibitors (e.g. Rlim) and/or inhibition of Myc activators (e.g. Aurka). This is borne out by the analysis of Myc co-factors in the Id and Id Robo1 KD RNA Seq data (Supplementary Table S5). Further work is needed to determine whether, and which, Myc cofactors are epistatic to Id-Robo1 signalling. Our data provides further evidence that Robo1 is an important suppressor of proliferation and self-renewal in TNBC and future work includes extending this work to models of human TNBC. Prior work showing high Robo1 expression association with good outcome in breast cancer is consistent with our finding (Chang et al., 2012). There has been substantial interest in targeting Myc (Shen et al., 2015; Yang et al., 2017) and Id1, but until now has been very challenging (Dang et al., 2017; Fong et al., 2003). We show that Id1 is able to reprogram Myc activity possibly via Robo1 and may provide an alternative strategy to target Myc-dependent transcription.

## Conclusion

We have demonstrated that breast cancer cells marked by Id expression have high propensity for key CSC phenotypes like proliferation and metastasis. We have uncovered a set of genes that are potential Id targets leading to identification of a mechanism which involves the negative transcriptional regulation of Robo1 by Id. This suggests an association between Id and Robo1 that correlates to the activation of a c-Myc driven proliferative and self-renewal program. Our observations suggest that we could exploit this pathway to target CSCs in the difficult to treat TNBC subtype.

## Abbreviations

CSC: Cancer stem cell
TNBC: Triple Negative Breast Cancer
Id: Id1 and Id3

## Availability of supporting data

All microarray and RNA Sequencing data has been deposited into the GEO database. Microarray (GSE129790) and RNA-Seq (GSE129858) datasets are available in SuperSeries GSE129859.

## Acknowledgements

We would like to thank the following people for their assistance with this manuscript: Ms. Nicola Foreman and Ms. Breanna Fitzpatrick for animal handling; Ms. Alice Boulghourjian and Ms. Anaiis Zaratzian for tissue processing and IHC staining; Mr. Rob Salomon and Mr. David Snowden for their help with flow sorting.

## Author’s contributions

RN and AS contributed to the conceptualizaton. WST, HH ASC, KH and SVK contributed to the methodology. DLR and CLC to the genomic analysis. RM, BAV, APT, AM, SJ, JY, IN, JSS and LAB contributed to the investigations. EKAM and SAO carried out the analysis on patient samples. ASM contributed to the resources. RN and NK wrote the original draft of the manuscript. MJN, CJO, SRL and WK reviewed, and edited the manuscript. NK contributed to the visualization. AS and RN contributed to the funding acquisition. All authors read and approved the final manuscript.

## Ethics approval

All experiments involving animal work were performed in accordance with the rules and regulations stated by the Garvan Institute Animal Ethics Committee.

## Supplementary information

Supplementary information is available as Supplementary Figures and Supplementary Tables.

