## Supplementary Information for "Id proteins promote a cancer stem cell phenotype in triple negative breast cancer via negative regulation of Robo1" (from another version of this preprint: version 4, 2020-03-10)

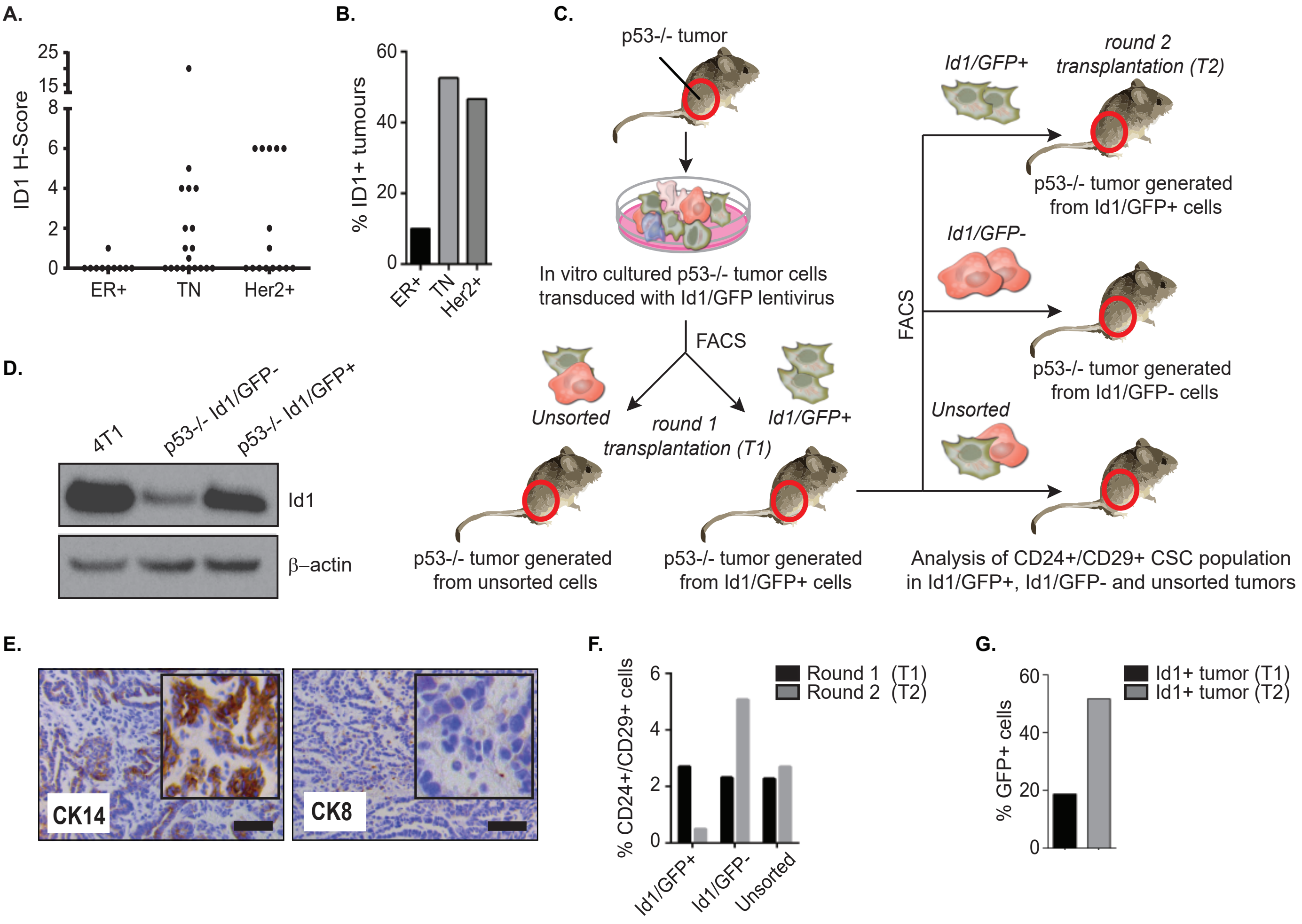


**Supplementary Figure S1. Identifying Triple Negative breast cancer models to dissect Id mechanism**.

**(A, B)** Quantification of ID1 expression in ER+, TN, and Her2-enriched breast cancer specimens. H score was determined by multiplying the percentage of ID1 positive tumour cells by the staining intensity (graded on a scale of 0-3). **(C)** Schematic showing the infection of p53-/- cultured tumour cells with the Id1/GFP reporter, followed by two transplantation rounds to determine the tumour initiating frequency of the Id1$^{+}$ versus Id1$^{-}$ tumour cells. **(D)** p53-/- tumour cells were transfected with the Id1/GFP reporter where 1.2kb of the Id1 proximal promoter has been placed upstream of GFP cDNA. Western Blot analysis demonstrate that FACS purified p53-/-/GFP$^{+}$ cells expressed high level of Id1 compared to the p53-/-/GFP$^{-}$ population. The 4T1 mouse mammary carcinoma cell line was used as a positive control for Id1 protein expression. **(E)** Representative IHC images of CK14 and CK8 expression in the Id1C3-Tag model, confirming its suitability as a model system. Bars = 50μM. **(F)** Comparison of CD29$^{+}$/CD24$^{+}$ cells (%) between groups gated for GFP expression in the p53-/- model. T1 and T2 indicate transplantation round 1 and 2, respectively. **(G)** The % of GFP $^{+}$ cells was calculated in the p53-/- tumours generated from Id1/GFP+ cells in round 1 and 2 transplantations. The % of GFP $^{+}$ cells was significantly higher in the second transplantation round.

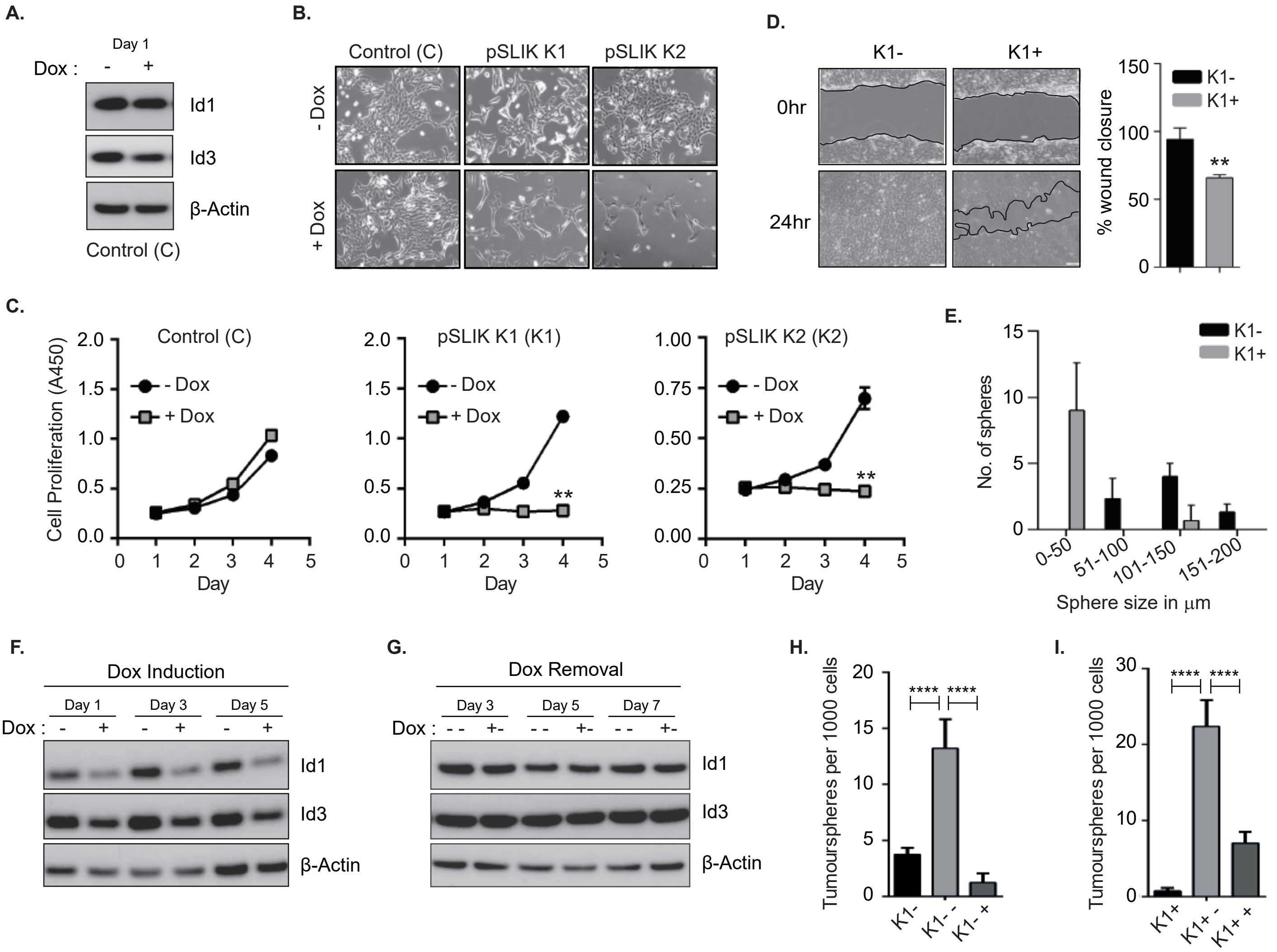


**Supplementary Figure S2. Characterization of the 4T1 knock down model system**.

**(A)** 4T1 pSLIK Control (C) cells were treated with Dox for 24 hours. Western analysis was performed on cell lysates harvested on day 1 post using anti-Id1 and Id3 antibodies and β-actin as loading control**.**
**(B)** Representative images of 4T1 pSLIK control (C) and 4T1 pSLIK shId clones (K1, K2) ± Dox.
**(C)** C, K1, K2 ± Dox cells were assayed for their proliferative potential. ** $p$<0.01 when Dox treated and untreated samples were compared. **(D)** Scratch-wound assay in K1± Dox showed a significant reduction in the migration of Id KD cells. **(E)** Quantification for the sizes of tumourspheres in control and Id KD cells.
**(F, G)** Re-expression of Id proteins following Dox removal in the 4T1 pSLIK shId polyclonal cells. 4T1 pSLIK shId pool cells were treated with Dox (Dox Induction), Dox was then omitted (Dox Removal, + -) from the medium for 7 days. Western analysis was performed on cell lysates harvested on day 3, 5 and 7 post Dox removal using anti-Id1 and Id3 antibodies and β-actin as loading control. **(H)** Primary tumour-spheres, not treated with Dox, were passaged to secondary tumourspheres in the absence or presence of Dox (-, - -, - +). Secondary tumourspheres were significantly less when the cells were depleted of Id (- +). Data are means ± SD (n=3) (**** $p$< 0.0001; One-way ANOVA). **(I)** Primary tumourspheres, previously treated with Dox (+) and depleted of Id expression, were passaged to secondary tumourspheres in the absence of Dox to allow re-expression of Id (+ -). Secondary tumourspheres from K1 cells cultured without Dox re-established their self-renewal potential by resuming the ability to form tumourspheres. Data are means ± SD (n=3) (**** $p$< 0.0001; One-way ANOVA).

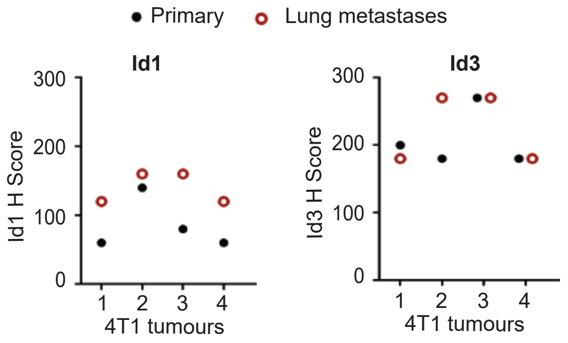


**Supplementary Figure S3. Id1 controls metastasis in TNBC.**
Graph showing the comparison of Id1 and Id3 expression in 4T1 primary tumours versus matched lung metastases (with and without Id KD). All cases showed an enrichment of Id1 expression in the lung metastasis. H score was determined by multiplying the percentage of Id1 or Id3 positive tumour cells by the staining intensity (graded on a scale of 0-3).

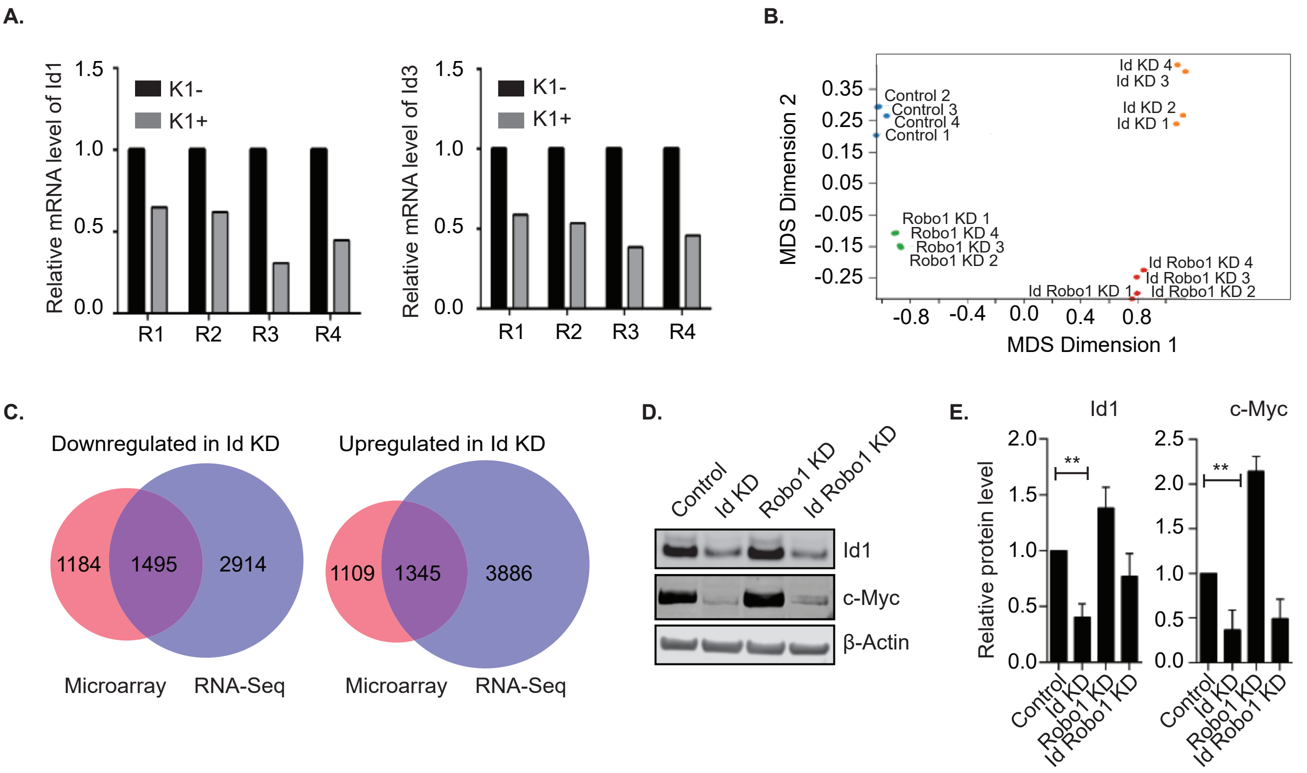


**Supplementary Figure S4. Gene expression analysis reveals a Myc signature**.

**(A)** Matching RNA samples (R1, R2, R3, R4) were analysed by quantitative real-time PCR for the expression of Id1 and Id3 genes using TaqMan® probed-based system. Gapdh was used as an endogenous control for normalisation of samples for any possible variations in amount of RNA added to each cDNA reaction as well as variation in PCR amplification efficiency. Relative mRNA level was obtained by comparing mRNA level in cells treated with Dox to the untreated cells which was set at 1. **(B)** RNA- Sequencing was performed on four replicates per condition and the MDS plots of the samples showed that the replicates cluster together. **(C)** Overlap of up and down regulated genes from the Microarray and RNA Seq Id1KD data. **(D)** Western blots showing the protein expression of Id1 and c-Myc in Control, Id KD, Robo1 KD and Id Robo1 KD cells. **(E)** Quantification for the protein expression of Id1 and c-Myc in Control, Id KD, Robo1 KD and Id Robo1 KD cells. β-Actin was used as the loading control and the expression levels of Id1 and c-Myc were normalized to that of β-Actin.

**Supplementary Table S1.** Top 25 up and down regulated genes identified in Id-depleted K1 cells ranked based on Q-value. Column 1 shows the gene symbols and annotation. Fold change in column 2 represents the ratio of gene expression change in Id1/3 knockdown versus control. Column 3 represents the Q-value for the difference between knockdown and control group. Column 4 shows the direction of change in gene expression.

| Gene | Fold Change | Q-value | Direction |
|---|---|---|---|
| Mx2 :: myxovirus (influenza virus) resistance 2 | 26.6015 | 7.59E-08 | UP |
| Oas1g :: 2'-5' oligoadenylatesynthetase 1G | 14.9574 | 1.04E-07 | UP |
| Oas3 :: 2'-5' oligoadenylatesynthetase 3 | 15.124 | 1.95E-07 | UP |
| Cmpk2 :: cytidine monophosphate (UMP-CMP) kinase 2, mitochondrial | 24.33 | 2.32E-07 | UP |
| Stat1 :: signal transducer and activator of transcription 1 | 6.8186 | 3.29E-07 | UP |
| Xaf1 :: XIAP associated factor 1 | 9.0697 | 4.06E-07 | UP |
| Usp18 :: ubiquitin specific peptidase 18 | 30.489 | 4.10E-07 | UP |
| Oas2 :: 2'-5' oligoadenylatesynthetase 2 | 36.1304 | 4.10E-07 | UP |
| Ifit1 :: interferon-induced protein with tetratricopeptide repeats 1 | 18.5318 | 4.58E-07 | UP |
| Gpr56 :: G protein-coupled receptor 56 | 21.8355 | 4.72E-07 | UP |
| Zbp1 :: Z-DNA binding protein 1 | 13.582 | 4.72E-07 | UP |
| Olfr65 :: olfactory receptor 65 | 8.3323 | 4.72E-07 | UP |
| Parp14 :: poly (ADP-ribose) polymerase family, member 14 | 5.9386 | 5.22E-07 | UP |
| Angptl4 :: angiopoietin-like 4 | 12.113 | 5.22E-07 | UP |
| Irf7 :: interferon regulatory factor 7 | 13.4442 | 5.22E-07 | UP |
| Gbp3 :: guanylate binding protein 3 | 10.1348 | 5.68E-07 | UP |
| Stat2 :: signal transducer and activator of transcription 2 | 5.0425 | 6.80E-07 | UP |
| Oasl2 :: 2'-5' oligoadenylatesynthetase-like 2 | 5.7617 | 7.46E-07 | UP |
| Bst2 :: bone marrow stromal cell antigen 2 | 8.4297 | 7.46E-07 | UP |
| Lypd3 :: Ly6/Plaur domain containing 3 | 4.2202 | 7.46E-07 | UP |
| Iigp1 :: interferon inducible GTPase 1 | 14.3364 | 7.93E-07 | UP |
| Pvrl1 :: poliovirus receptor-related 1 | 4.7014 | 7.93E-07 | UP |
| Oas1b :: 2'-5' oligoadenylatesynthetase 1B | 9.7557 | 8.90E-07 | UP |
| Megf10 :: multiple EGF-like-domains 10 | 11.0025 | 1.04E-06 | UP |
| Rtp4 :: receptor transporter protein 4 | 9.0028 | 1.04E-06 | UP |
| H19 :: H19 fetal liver mRNA | 2.8526 | 3.68E-06 | DOWN |
| Mpzl2 :: myelin protein zero-like 2 | 3.1982 | 3.86E-06 | DOWN |
| Cpox :: coproporphyrinogen oxidase | 2.5431 | 4.32E-06 | DOWN |
| Gpr116 :: G protein-coupled receptor 116 | 2.9257 | 5.34E-06 | DOWN |
| Cep78 :: centrosomal protein 78 | 2.5606 | 8.09E-06 | DOWN |
| Gpt2 :: glutamic pyruvate transaminase (alanine aminotransferase) 2 | 3.2592 | 8.29E-06 | DOWN |
| Cth :: cystathionase (cystathionine gamma-lyase) | 4.9146 | 1.01E-05 | DOWN |
| Zfp420 :: zinc finger protein 420 | 2.6201 | 1.07E-05 | DOWN |

| | | | |
|---|---|---|---|
| Gja1 :: gap junction protein, alpha 1 | 2.2993 | 1.73E-05 | DOWN |
| Cldn9 :: claudin 9 | 4.2349 | 1.82E-05 | DOWN |
| Rtkn2 :: rhotekin 2 | 3.3638 | 1.93E-05 | DOWN |
| Aqp1 :: aquaporin 1 | 3.4829 | 2.80E-05 | DOWN |
| Fermt1 :: fermitin family homolog 1 (Drosophila) | 2.1462 | 3.05E-05 | DOWN |
| Taf5 :: TAF5 RNA polymerase II, TATA box binding protein (TBP)-associated factor | 1.9337 | 3.62E-05 | DOWN |
| Slc7a11 :: solute carrier family 7 (cationic amino acid transporter, y+ system), member 11 | 7.3157 | 3.72E-05 | DOWN |
| B4galnt4 :: beta-1,4-N-acetyl-galactosaminyl transferase 4 | 2.2432 | 3.81E-05 | DOWN |
| Gls2 :: glutaminase 2 (liver, mitochondrial) | 2.3353 | 3.93E-05 | DOWN |
| Fbln2 :: fibulin 2 | 2.0405 | 4.05E-05 | DOWN |
| 2610021K21Rik :: RIKEN cDNA 2610021K21 gene | 2.9111 | 4.51E-05 | DOWN |
| Gstcd :: glutathione S-transferase, C-terminal domain containing | 1.8478 | 0.0000465 | DOWN |
| Rps6ka6 :: ribosomal protein S6 kinase polypeptide 6 | 2.1709 | 5.28E-05 | DOWN |
| Hspa1l :: heat shock protein 1-like | 1.9999 | 5.89E-05 | DOWN |
| Dyrk3 :: dual-specificity tyrosine-(Y)-phosphorylation regulated kinase 3 | 2.8598 | 6.15E-05 | DOWN |
| Hmgn5 :: high-mobility group nucleosome binding domain 5 | 3.2189 | 6.36E-05 | DOWN |
| Deptor :: DEP domain containing MTOR-interacting protein | 4.1093 | 6.37E-05 | DOWN |

**Supplementary Table S2.** Top 20 gene sets identified from the C5 GO gene sets that are enriched in the Id- knockdown K1 cells.

| C5 GO Gene Set | Normalised Enrichment Score | P value | Direction |
|---|---|---|---|
| IMMUNE_RESPONSE | 2.136929 | <0.0001 | UP |
| G_PROTEIN_COUPLED_RECEPTOR_BINDING | 2.1243143 | <0.0001 | UP |
| CHEMOKINE_RECEPTOR_BINDING | 2.111146 | <0.0001 | UP |
| CHEMOKINE_ACTIVITY | 2.0914385 | <0.0001 | UP |
| I_KAPPAB_KINASE_NF_KAPPAB_CASCADE | 2.0558612 | <0.0001 | UP |
| DEFENSE_RESPONSE | 2.0529644 | <0.0001 | UP |
| JAK_STAT_CASCADE | 2.0520313 | <0.0001 | UP |
| HEMATOPOIETIN_INTERFERON_CLASSD200_DOMAIN_CYTOKINE_RECEPTOR_ACTIVITY | 2.0426128 | <0.0001 | UP |
| LOCOMOTORY_BEHAVIOR | 2.0126882 | <0.0001 | UP |
| REGULATION_OF_I_KAPPAB_KINASE_NF_KAPPAB_CASCADE | 2.0115595 | <0.0001 | UP |
| M_PHASE | -2.5211284 | <0.0001 | DOWN |
| CELL_CYCLE_PROCESS | -2.5124967 | <0.0001 | DOWN |
| RNA_PROCESSING | -2.4857798 | <0.0001 | DOWN |
| CHROMOSOME | -2.3979888 | <0.0001 | DOWN |
| M_PHASE_OF_MITOTIC_CELL_CYCLE | -2.3873682 | <0.0001 | DOWN |
| MITOSIS | -2.386327 | <0.0001 | DOWN |
| CHROMOSOMEPERICENTRIC_REGION | -2.366616 | <0.0001 | DOWN |
| SPINDLE | -2.3466449 | <0.0001 | DOWN |
| CHROMOSOME_ORGANIZATION_AND_BIOGENESIS | -2.3239574 | <0.0001 | DOWN |
| MITOTIC_CELL_CYCLE | -2.3222191 | <0.0001 | DOWN |

**Supplementary Table S3.** Top 20 gene sets identified from the C6 Oncogenic Signatures that are enriched in the Id- knockdown K1 cells

| C6 Oncogenic Gene Set | Normalised Enrichment Score | P value | Direction |
|---|---|---|---|
| LTE2_UP.V1_DN | 2.389604 | <0.0001 | UP |
| MEK_UP.V1_DN | 2.244798 | <0.0001 | UP |
| VEGF_A_UP.V1_UP | 2.124735 | <0.0001 | UP |
| WNT_UP.V1_DN | 2.070847 | <0.0001 | UP |
| PKCA_DN.V1_UP | 1.969476 | <0.0001 | UP |
| MYC_UP.V1_DN | 1.865835 | <0.0001 | UP |
| BMI1_DN_MEL18_DN.V1_DN | 1.860512 | <0.0001 | UP |
| MEL18_DN.V1_DN | 1.845022 | <0.0001 | UP |
| BMI1_DN.V1_DN | 1.83551 | <0.0001 | UP |
| STK33_UP | 1.819187 | <0.0001 | UP |
| RPS14_DN.V1_DN | -2.3749187 | <0.0001 | DOWN |
| HOXA9_DN.V1_DN | -2.214685 | <0.0001 | DOWN |
| CSR_LATE_UP.V1_UP | -2.1530662 | <0.0001 | DOWN |
| PRC2_EZH2_UP.V1_UP | -2.0872 | <0.0001 | DOWN |
| VEGF_A_UP.V1_DN | -2.0221975 | <0.0001 | DOWN |
| RB_P107_DN.V1_UP | -2.0133445 | <0.0001 | DOWN |
| E2F1_UP.V1_UP | -2.0126765 | <0.0001 | DOWN |
| MYC_UP.V1_UP | -1.8403969 | <0.0001 | DOWN |
| STK33_DN | -1.8023152 | <0.0001 | DOWN |
| NFE2L2.V2 | -1.7664328 | <0.0001 | DOWN |

**Supplementary Table S4.** Top 20 gene sets identified from the C2 curated gene sets that are enriched in the Id-knockdown K1 cells.

| C2 Curated Gene Set | Normalised Enrichment Score | P value | Direction |
|---|---|---|---|
| BROWNE_INTERFERON_RESPONSIVE_GENES | 2.893714 | <0.0001 | UP |
| TAKEDA_TARGETS_OF_NUP98_HOXA9_FUSION_3D_UP | 2.889862 | <0.0001 | UP |
| ICHIBA_GRAFT_VERSUS_HOST_DISEASE_D7_UP | 2.805356 | <0.0001 | UP |
| SANA_TNF_SIGNALING_UP | 2.760641 | <0.0001 | UP |
| SANA_RESPONSE_TO_IFNG_UP | 2.652481 | <0.0001 | UP |
| DER_IFN_ALPHA_RESPONSE_UP | 2.650679 | <0.0001 | UP |
| DAUER_STAT3_TARGETS_DN | 2.568591 | <0.0001 | UP |
| DER_IFN_BETA_RESPONSE_UP | 2.544266 | <0.0001 | UP |
| MOSERLE_IFNA_RESPONSE | 2.528648 | <0.0001 | UP |
| RADAEVA_RESPONSE_TO_IFNA1_UP | 2.456697 | <0.0001 | UP |
| ROSTY_CERVICAL_CANCER_PROLIFERATION_CLUSTER | -3.2496483 | <0.0001 | DOWN |
| SOTIRIOU_BREAST_CANCER_GRADE_1_VS_3_UP | -3.1307108 | <0.0001 | DOWN |
| WONG_EMBRYONIC_STEM_CELL_CORE | -3.0442135 | <0.0001 | DOWN |
| GRAHAM_CML_DIVIDING_VS_NORMAL_QUIESCENT_UP | -2.9951138 | <0.0001 | DOWN |
| KOBAYASHI_EGFR_SIGNALING_24HR_DN | -2.950584 | <0.0001 | DOWN |
| LEE_EARLY_T_LYMPHOCYTE_UP | -2.9474769 | <0.0001 | DOWN |
| PUJANA_BRCA2_PCC_NETWORK | -2.91798 | <0.0001 | DOWN |
| LI_WILMS_TUMOR_VS_FETAL_KIDNEY_1_DN | -2.8696544 | <0.0001 | DOWN |
| SHEDDEN_LUNG_CANCER_POOR_SURVIVAL_A6 | -2.8691382 | <0.0001 | DOWN |
| FURUKAWA_DUSP6_TARGETS_PCI35_DN | -2.841147 | <0.0001 | DOWN |

**Supplementary Table S5.** List of differentially expressed Myc co-factors in Id + Robo1 KD in comparison to Id1/3 KD alone obtained from the RNA-Seq data. Red indicates negative regulators and green indicates positive regulators.

| Downregulated by Robo1 KD in the absence of Id1/3 | Upregulated by Robo1 KD in the absence of Id1/3 | Unchanged by Robo1 KD in the absence of Id1/3 |
|---|---|---|
| Bptf | Nme2 | Ash2l |
| Pim1 | MlxIP | Bptf |
| Rlim | Mlx | Brd4 |
| Zbtb17 | Actl6a | Cdk8 |
| Smad3 | Kat2a | Cdk9 |
| Hbp1 | Ruvbl2 | Dot1l |
| Mxd4 | Skp2 | Ep300 |
| Tsc22d1 | Banf1 | Ep400 |
| Hdac2 | Ash2l | Kat5 |
| Mycl | Atad2 | Kdm5a |
| Rnf115 | Ccnt1 | Max |
| | Npm1 | Med26 |
| | Aurka | Myc |
| | Clock | Mycn |
| | Mxd3 | Sirt1 |
| | Hdac1 | Smarcb1 |
| | Suds3 | Snip1 |
| | Sap30 | Trrap |
| | Sap18 | Arntl |
| | Rbbp7 | Fbxw7 |
| | Fbxw7 | Huwe1 |
| | Trpc4ap | Mga |
| | Bin1 | Mnt |
| | Rbl1 | Mxd1 |
| | | Rpl11 |
| | | Sin3a |
| | | Sin3b |
| | | Btg1 |

**Supplementary Table S6.** The list and the concentrations of antibodies used for western blotting in the study.

| Antibody | Manufacturer | Catalogue # | Concentration |
|---|---|---|---|
| Rabbit anti-mouse Id1 (primary) | BioCheck (Burlingame, CA, USA) | BCH-1/37-2 | 1:500 |
| Rabbit anti-human/mouse Id3 (primary) | BioCheck (Burlingame, CA, USA) | BCH-4/#17-3 | 1:500 |
| Mouse anti- β-actin (primary) | AbCam (Cambridge, UK) | Ab6276 | 1:4000 |
| Horse anti-mouse IgG-HRP (secondary) | Cell Signaling (Beverly, MA, USA) | 7054 | 1:8000 |
| Goat anti-rabbit IgG-HRP (secondary) | Cell Signaling (Beverly, MA, USA) | 7054 | 1:5000 |

**Supplementary Table S7.** The list and the concentrations of antibodies used for Immunohistochemistry in the study.

| Antibody | Manufacturer | Catalogue # | Concentration |
|---|---|---|---|
| Rabbit anti-mouse Id1 (primary) | BioCheck (Burlingame, CA, USA) | BCH-1/37-2 | 1:50 |
| Rabbit anti-human/mouse Id3 (primary) | BioCheck (Burlingame, CA, USA) | BCH-4/#17-3 | 1:50 |
| Rabbit anti-human Id1 (primary) | BioCheck (Burlingame, CA, USA) | BCH-1/195-14 | 1:50 |
| Rabbit anti-human/mouse CK14 (Primary) | Covance (Princeton, NJ, USA) | PRB-155P | 1:8000 |
| Rabbit anti-human/mouse CK8 (Primary) | Cell Signaling (Beverly, MA, USA) | 7054 | 1:5000 |
